# Ecology, not host phylogeny, shapes the oral microbiome in closely related species of gorillas

**DOI:** 10.1101/2022.06.06.494923

**Authors:** Markella Moraitou, Adrian Forsythe, James A. Fellows Yates, Jaelle C. Brealey, Christina Warinner, Katerina Guschanski

**Affiliations:** Animal Ecology, Department of Ecology and Genetics, Uppsala University, 75236 Uppsala, Sweden; Department of Archaeogenetics, Max Planck Institute for Evolutionary Anthropology, 04103 Leipzig, Germany; Department of Paleobiotechnology, Leibniz Institute for Natural Product Research and Infection Biology Hans Knöll Institute, 07745 Jena, Germany; Department of Natural History, NTNU University Museum, Norwegian University of Science and Technology, 7491 Trondheim, Norway; Faculty of Biological Sciences, Friedrich Schiller University, 07743 Jena, Germany; Department of Anthropology, Harvard University, Cambridge, MA 02138; Science for Life Laboratory, 75237 Uppsala, Sweden; Institute of Evolutionary Biology, School of Biological Sciences, University of Edinburgh, Edinburgh EH9 3FL, United Kingdom

**Keywords:** ancient DNA, dental calculus, museum specimens, gorilla, metagenome-assembled genomes, shotgun metagenomics

## Abstract

Host-associated microbiomes are essential for a multitude of biological processes. Placed at the contact zone between external and internal environments, the little-studied oral microbiome has important roles in host physiology and health. Here we investigate the contribution of host evolutionary relationships and ecology in shaping the oral microbiome in three closely related gorilla subspecies (mountain, Grauer’s, and western lowland gorillas) using shotgun metagenomics of 46 museum-preserved dental calculus samples. We find that the oral microbiomes of mountain gorillas are functionally and taxonomically distinct from the other two subspecies, despite close evolutionary relationships and geographic proximity with Grauer’s gorillas. Grauer’s gorillas show intermediate bacterial taxonomic and functional, and dietary profiles. Altitudinal differences in gorilla subspecies ranges appear to explain these patterns, proposing a close connection between dental calculus microbiome and the environment, which is further supported by the presence of gorilla subspecies-specific phyllosphere/rhizosphere taxa. Mountain gorillas show high abundance of nitrate-reducing oral taxa, which may contribute to high altitude adaptation by modulating blood pressure. Our results suggest that ecology, rather than evolutionary relationships and geographic proximity, primarily shape the oral microbiome in these closely related species.

## Background

The microbial communities that are found on and inside multicellular organisms are not only remarkably diverse, but also play a crucial role in a vast set of biological processes, such as energy uptake (Arora and Sharma, 2011; Nieuwdorp et al., 2014), detoxification (Nichols et al., 2019; Turner and Bucking, 2019), immune responses (Nikitakis et al., 2017; West et al., 2015) and even neurochemical and hormonal processes that eventually influence behaviour (Suzuki, 2017; Vuong et al., 2017). The extensively studied gut microbiome is shaped by a multitude of factors, including host phylogenetic relationships, social interactions, and dietary adaptations (Moeller et al., 2016; Nishida and Ochman, 2018; Youngblut et al., 2019). In comparison to the gut microbiome, the oral microbiome has received far less attention. Yet, it is of particular importance, as it connects the external and the internal environments and is located directly at the entry point to the digestive and the respiratory tracts. The oral microbiome plays an important role in oral diseases, such as caries and periodontitis, and has been implicated in systemic disorders including cardiovascular disease (Nakano et al., 2011), atherosclerosis (Teles and Wang, 2011), Alzheimer’s disease (Olsen and Singhrao, 2015), several cancers (Flynn et al., 2016; Gao et al., 2016) and preterm births (Cobb et al., 2017).

Although studies have linked oral microbiome composition to both dietary habits (Adler et al., 2016; Hyde et al., 2014b; Janiak et al., 2021; Nearing et al., 2020) and host evolutionary relationships (Brealey et al., 2020; Ozga and Ottoni, 2021), it is unclear which of the two predominantly drives the evolution of the oral microbiome and how they interact. Early studies suggested that the oral microbiome is strongly heritable and mostly transferred vertically (Corby et al., 2007; Li et al., 2007), with neonate microbiomes resembling those of their mothers (Dominguez-Bello et al., 2010). Moreover, studies in wild animals have indicated that different species harbour distinct oral microbiota (Brealey et al., 2020; Ozga and Ottoni, 2021). Both observations suggest that host evolutionary relationships have a strong effect on oral microbiome structure. Other studies have highlighted the effect of diet on the oral microbiome (Janiak et al., 2021; Wade, 2013), for instance by uncovering differences between young infants and adults of humans and non-human primates, hence providing a potential link with age-related dietary changes (Cephas et al., 2011). However, distinguishing between host evolutionary and ecological factors as well as their contribution to oral microbiome evolution remains difficult, as most previous studies have focused on rather distantly related host species that occupy distinct ecological niches (Boehlke et al., 2020; Li et al., 2013; Smith et al., 2021; Soares-Castro et al., 2019).

In recent years, dental calculus - the calcified form of dental plaque that forms on the teeth of mammals - has emerged as a useful material for the study of oral microbiomes across diverse mammalian species (Brealey et al., 2020; Fellows Yates et al., 2021; Ottoni et al., 2019; Warinner et al., 2014). Since dental plaque undergoes periodic mineralizations, it effectively fossilises on the living host, reducing postmortem contamination from environmental microorganisms (Warinner et al., 2015). It thus preserves a snapshot of oral and respiratory microbial communities (Warinner et al., 2014), dietary components (Adler et al., 2013; Radini et al., 2017; Sawafuji et al., 2020) and host DNA (Mann et al., 2018). Museum collections can be efficiently used for the study of oral microbiomes from wild animals (Brealey et al., 2020; Fellows Yates et al., 2021), minimising the exposure and disturbances associated with sampling from live hosts. Furthermore, the detection of damage patterns, a common verification method for ancient DNA (Briggs et al., 2007), can also be implemented on such historical specimens to distinguish endogenous taxa from modern contaminants (Fellows Yates et al., 2021). Previous research suggests that microbial communities in historical and modern dental calculus are remarkably similar (Velsko et al., 2019), therefore studying museum specimens provides reliable information about present-day microbiomes. The exceptional preservation in the calcified matrix also allows assembly of near complete metagenome-assembled genomes (MAGs) (Brealey et al., 2020).

In this study, we use dental calculus to uncover ecological and evolutionary factors that drive oral microbiome evolution in a group of closely related species. We focus on gorillas, which consist of two species, western gorilla (*Gorilla gorilla*) and eastern gorilla (*G. beringei*), each of which is further divided into two subspecies. Our sampling covers three of the four subspecies: western lowland gorilla (*G. g. gorilla*), Grauer’s gorilla (*G. b. graueri*) and mountain gorilla (*G. b. beringei*). Eastern and western gorillas diverged approximately 250,000 years ago, with evidence for substantial gene flow until more recently (McManus et al., 2015; Xue et al., 2015). Western lowland gorillas are found in western equatorial Africa, primarily at low elevations below 500 m above sea level (masl). Their diet consists mainly of various plant parts and fruits (Rogers et al., 2004). The eastern subspecies - mountain and Grauer’s gorillas - occur on the eastern side of the Congo basin. They are estimated to have diverged from each other 10,000 years ago (Roy et al., 2014). Grauer’s gorillas occupy the largest altitudinal range of all gorillas, ranging from 500 to 2,900 masl (Plumptre et al., 2016). Mountain gorillas are found at even higher elevations, with most of the range of the Virunga Massif population above 2,200 masl and extending to up to 3,800 masl. Consequently, these altitudinal differences lead to different dietary profiles among populations and subspecies (Doran et al., 2002; Ganas et al., 2004; Michel et al., 2022; Rogers et al., 2004; van der Hoek et al., 2021), primarily reflecting availability of fruits that are highly seasonal and become scarce with increasing altitude (Ganas et al., 2004).

The presence of ecological and dietary differences in these closely related gorilla subspecies provides an opportunity to evaluate the effects of host evolutionary relationships and ecological factors on oral microbiomes. We used shotgun metagenomic sequencing of dental calculus samples from museum specimens (collected between 1910 and 1986) to taxonomically and functionally characterise the oral microbiota, as well as to assemble bacterial MAGs and identify dietary components from the three gorilla subspecies. Between closely related gorilla subspecies, ecology has a large impact on the taxonomic composition and function capacities of the oral microbiome. Our analyses highlight subspecies-specific differences that can be attributed ecological/dietary differences, and bacterial members of the oral microbiome that may contribute to adaptation to high-altitude lifestyle.

## Results

### Data pre-processing and confirmation of oral microbial signature

We produced paired-end shotgun sequencing data from the gorilla dental calculus of 26 newly collected samples, and complemented these with an additional 31 dental calculus samples from previously published studies (Brealey et al., 2020; Fellows Yates et al., 2021) (Table S1). Four dental calculus samples were sequenced independently in this study and by Fellows Yates et al. (2021). At a later stage of the analysis, this set of technical duplicates allowed us to assess the effect of the dataset in our taxonomic analysis, but only the sample with the largest number of reads for each sample pair was retained for further analyses.

Raw sequencing data consisted of, on average, 15,533,319 reads (mean value) per sample (ranging from 1,032 to 95,367,058; Table S1) and 105,653 reads per negative control (both extraction and library preparation, ranging from 19 to 1,132,867; Table S1). After raw sequence pre-processing (which included removal of poly-G tails, adapter and barcode sequences, merging of forward and reverse reads, and quality filtering), *phiX* and gorilla/human sequences were removed by mapping against respective reference genomes and the resulting unmapped reads were used for taxonomic classification using Kraken2 (Wood et al., 2019) with the standard database, which includes all bacterial, archaeal and viral genomes from NCBI. Samples with low read counts, low proportion of oral taxa, as well as duplicate samples (those processed at two different facilities), were removed (Methods). The final dataset consisted of 46 dental calculus samples (13 western lowland gorillas, 17 mountain gorilla, 16 Grauer’s gorilla; Figure S1), each containing 561,978-77,307,443 reads (mean: 11,365,074, SD: 15,577,988; Table S1). We then applied a multi-step procedure, relying on negative controls and museum environmental samples, to remove contaminant taxa (Methods). The final dataset contained a total of 1007 microbial species (in 430 genera), of which 3.4% (n=34) were members of the core hominid oral microbiome (Fellows Yates et al., 2021) and 4.2% (n=42) members of the Human Oral Microbiome Database (HOMD; (Chen et al., 2010); Table S2). These microbial species accounted for ca. 14% of the total microbial non-normalised abundance.

The final dataset also included six genera that have been listed as common contaminants by Salter *et al*. (2014) and Weyrich *et al*. (2019), but contain known oral taxa (Chen et al., 2010; Fellows Yates et al., 2021), including *Streptococcus oralis* and *Staphylococcus saprophyticus.* Species belonging to these genera were retained after confirming, where possible, that they exhibit typical post-mortem DNA damage patterns (Methods).

### Partial mitochondrial genomes from dental calculus allow host subspecies identification

We recovered host mitochondrial sequences from 44 of 46 samples, which we used to confirm gorilla subspecies identity. On average, 31.2% (0.44 - 94.2%; Table S3) of the reference mitochondrial genome was covered by mapped reads, with notable variation in the proportion of genome covered and coverage depth (0.01 - 77.04%, average: 3.27%) between samples. Sample weight (information available for n=24 samples), sample age (n=36), or dataset (newly generated versus previously published by Fellows Yates et al. (2021)) (n=46) did not have an effect on mtDNA genome completeness (ANOVA, p>0.54), whereas host subspecies identity was marginally significant (ANOVA, F-value=3.12, p=0.054). Mountain gorillas had a larger proportion of the mtDNA genome covered by mapped reads than other gorilla subspecies. The cause of this difference is unknown but could be related to factors affecting the shedding of host DNA into the oral environment, such as differences in dietary abrasiveness, salivary flow, or inflammation.

To assist with host taxon assignment, we identified 403 diagnostic sites between western and eastern gorillas and 72 diagnostic sites between the two eastern gorilla subspecies (mountain and Grauer’s gorillas) using published gorilla mitochondrial genomes (Methods). In most samples, reads mapped to only a small number of diagnostic sites (Table S3). We accepted molecular taxon assignment for samples with reads mapping to at least six diagnostic sites in six separate reads. For six samples, data was insufficient to distinguish between eastern and western gorilla species and for additional six samples we could not reliably distinguish between mountain and Grauer’s gorilla subspecies. In these cases, we accepted the host subspecies identification based on museum records, after consulting collection locality, where possible. All but one sample containing a sufficient number of diagnostic sites were successfully assigned to their reported subspecies. The exception was sample MTM010 (museum accession: 631168, Swedish Museum of Natural History - NRM), which was confirmed to be a mountain gorilla, in accordance with the museum records, although it was previously reported as a western lowland gorilla based on preliminary genomic information (Fellows Yates et al., 2021).

### Oral microbiome diversity and composition differ by host subspecies

We evaluated the differences in microbial alpha diversity between gorilla subspecies by measuring species richness and community evenness (Figure 1). Host subspecies had only a marginal effect on the normalised microbial richness (ANOVA, p=0.061; Figure 1a) but a significant effect on the transformed community evenness (ANOVA, p<0.001; Figure 1b), with the microbial communities of mountain gorillas showing the lowest evenness.

**Figure 1.**
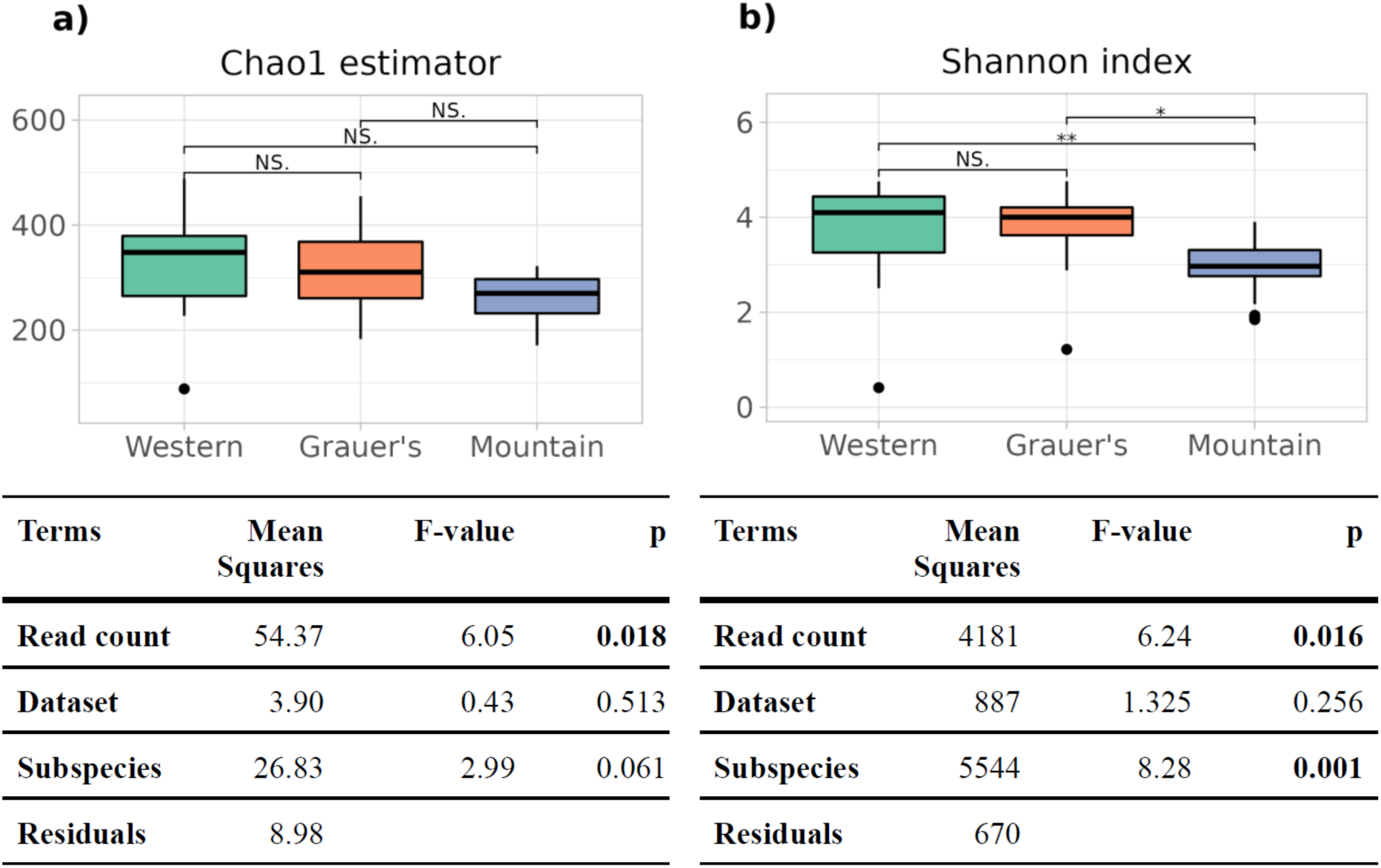
(a) Estimates of community richness, using Chao1 index and (b) community evenness, using Shannon index, alongside corresponding ANOVA results (tables). Pairwise comparisons in (a) and (b) show FDR-adjusted p-values of the Tukey’s test (NS=not significant, *p<0.05, **p<0.01). For the ANOVA models, sqrt(500-x) and exp(x) transformations were implemented on the Chao1 estimator and Shannon index values, respectively.

Microbiome composition differed by gorilla subspecies (Figure 2, Table 1). This effect persisted even after other potentially confounding factors, such as sequencing depth and dataset, were taken into account. Sequencing depth, included as read count, and dataset were significant factors for the presence-absence-based Jaccard measure, whereas neither of these factors significantly affected relative abundances as estimated with Aitchison distances (Table 1). We confirmed that belonging to different datasets had little to no effect on relative abundance by considering three pairs of samples that were derived from the same museum specimen, but were independently sequenced in this study and in Fellows Yates et al. (2021): the duplicate samples appeared close to each other on ordination plots, in particular on the plot based on Aitchison distances (Figure S2). The fourth sample pair (G0004-IBA002) was excluded, as both replicates had lower than 3% oral component, according to microbial source tracking using FEAST (Shenhav et al., 2019) (Figure S3). Pairwise PERMANOVAs (PERMutational ANalysis Of VAriance) (Anderson, 2001) showed strong differentiation between mountain gorillas and the other two subspecies, using both presence-absence and abundance metrics (Table 1). Lastly, we found no significant effect of sex on the oral microbiome using a subset of samples that could be genetically sexed (n=38; Methods; Table S4).

**Figure 2.**
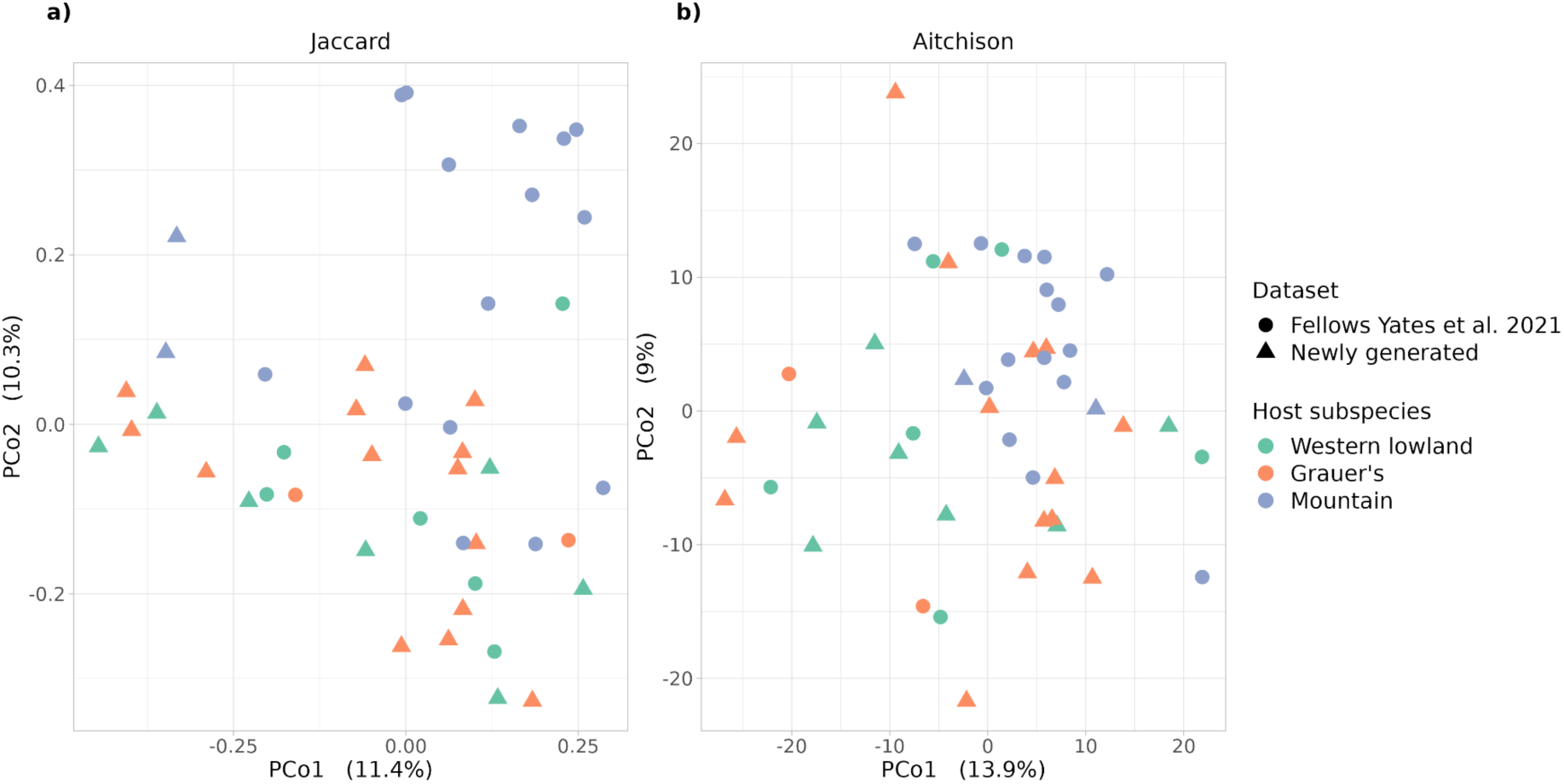
Principal coordinate analysis (PCoA) plots of the individual microbiomes based on a) Jaccard distance and b) Aitchison distance. Host subspecies is displayed in different colours, whereas the dataset is indicated by different shapes (circles for Fellows Yates et al. (2021) and triangles for the newly generated data).

**Table 1.** PERMANOVA results, showing the effect size (R^2^) and p-value of factors putatively contributing to the variance among sampled microbiota, using Jaccard (reflecting presence-absence of taxa) and Aitchison (reflecting relative abundance of taxa) distances. Pairwise PERMANOVA between subspecies show the effect size (R^2^) and p-values adjusted for false discovery rate. P-values below 0.05 are shown in bold.

|  |  | Jaccard distances |  | Aitchison distances |  |
| --- | --- | --- | --- | --- | --- |
| Terms | Within-factor comparison | $R^2$ | p | $R^2$ | p |
| Read count |  | 0.069 | < **0.001** | 0.023 | 0.297 |
| Dataset |  | 0.037 | **0.005** | 0.029 | 0.083 |
| Subspecies |  | 0.067 | < **0.001** | 0.074 | **0.001** |
|  | Western lowland vs Grauer's | 0.029 | 0.853 | 0.041 | 0.223 |
|  | Western Lowland vs Mountain | 0.086 | **0.001** | 0.085 | **0.001** |
|  | Grauer's vs Mountain | 0.073 | **0.001** | 0.072 | **0.001** |
| Residuals |  | 0.826 | - | 0.874 | - |

Among the 1007 microbial (species-level) taxa in our dataset, we detected 91 that significantly differed in abundance among gorilla subspecies and 11 that significantly differed in abundance between the datasets, using ANCOM (Mandal et al., 2015) (Table S5). Ten of these 11 taxa also differed in abundance by subspecies but because they could reflect dataset-specific artefacts, we removed the entire genera they belonged to from the list of subspecies-associated taxa. Among the remaining 78 differentially abundant taxa, all but six were absent from at least one host subspecies. The abundances of these taxa were comparable to not differentially abundant taxa (n=929) (t-test: t(123.59)=1.24, p=0.22): 37 taxa ranked above average and 75 were in the third quartile of abundance ranks, so that their absence from some host subspecies is unlikely to be explained by non-detection due to low abundance.

Mountain gorillas, in particular, appeared to be missing many of these differentially abundant microbial taxa (Figure 3) — which is consistent with the general observation of a marginally reduced microbial richness in this subspecies (Figure 1) — even after accounting for the effect of read depth. Microbial taxa that were absent in mountain gorillas included the orders *Rhodobacteriales*, *Pseudonocardiales*, *Corynebacteriales* (represented mainly by *Corynebacterium* and *Mycolicibacterium*), *Bacillales* (including *Staphylococcus*), and *Rhizobiales* (including bacteria associated with the *Fabaceae* rhizosphere, such as *Agrobacterium deltaense, A. fabacearum* (Delamuta et al., 2020; Yan et al., 2017), and three *Rhizobium* species (Poole et al., 2018)). The presence of rhizosphere- and phyllosphere-associated taxa in the other two subspecies, western lowland and Grauer’s gorillas, may reflect habitat or dietary differences among the subspecies. Microbial taxa enriched in mountain gorillas primarily belonged to the orders *Enterobacterales* and certain *Lactobacillales*, like *Streptococcus* sp. and *Lactobacillus gasseri*. The microbiomes of Grauer’s gorillas resembled those of western lowland gorillas, both overall (Figure 2, Table 1) and in terms of differentially abundant taxa (Figure 3). However, they also shared some similarities with mountain gorillas (e.g. a presence of *Limosilactobacillus*/*Lactobacillus* species, which were absent in western lowland gorillas), showing an intermediate or mixed composition (Figure 3).

**Figure 3.**
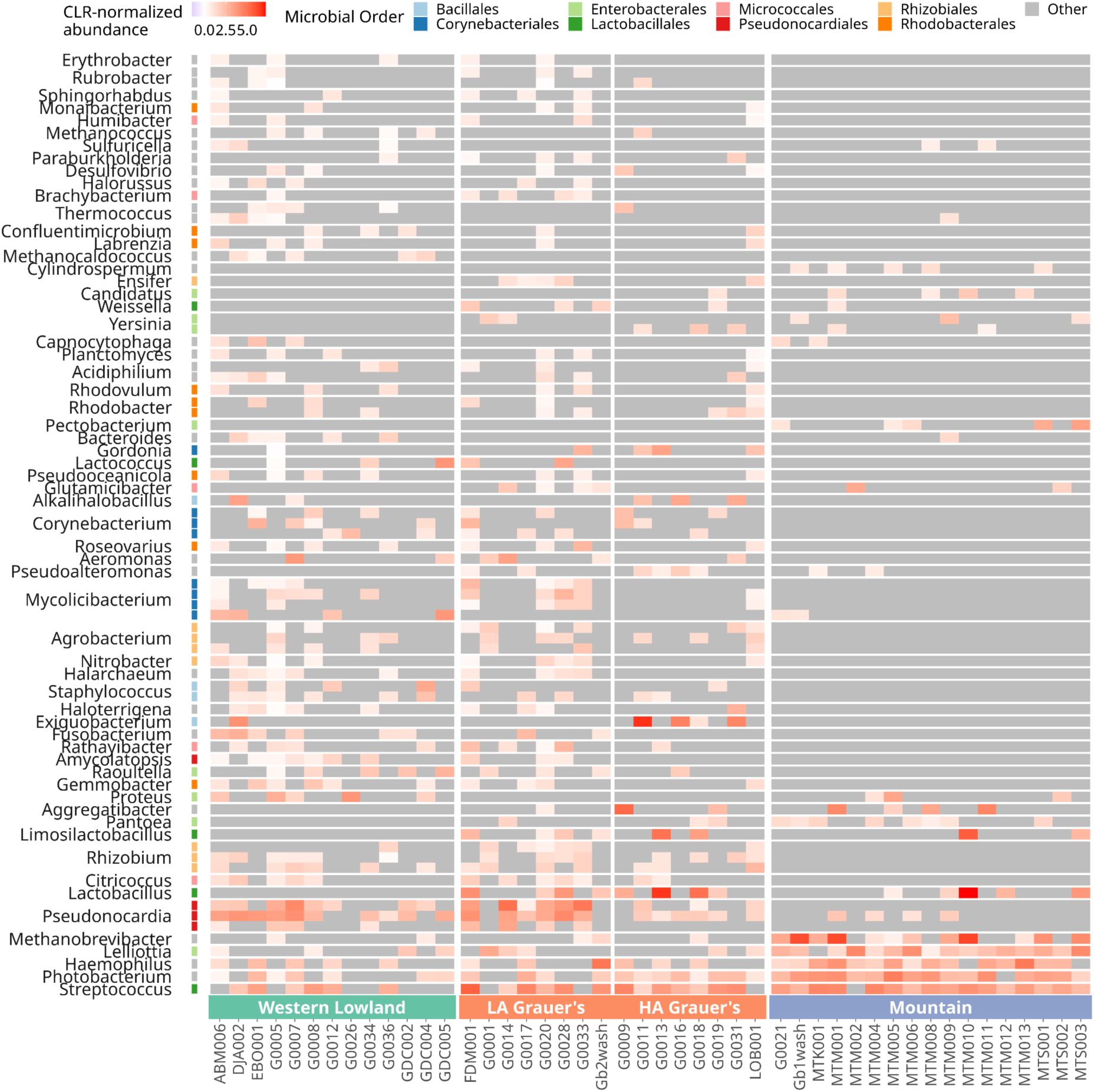
Heatmap depicting centred log-ratio (CLR) normalised abundances of 78 differentially abundant microbial taxa, coloured by taxonomic order (y axis), per sample (x axis). Due to the CLR normalisation, very low abundances or those equal to zero appear as negative values. For clarity, absent taxa (non-normalized abundance equal to zero) are shown in grey.

### Gorilla subspecies harbour functionally distinct oral communities

After removing reads mapping to identified contaminants, the retained (“decontaminated”) metagenomic reads were functionally characterised with an assembly-free approach using the HUMAnN2 pipeline (Franzosa et al., 2018) (Methods). The normalised abundances of gene families were regrouped under gene ontology (GO) terms, and were aggregated across microbial taxa. Results of functional analyses were broadly consistent with those observed for community-level taxonomy. Specifically, gorilla subspecies was marginally significant (p=0.06) in explaining functional differences in the oral microbiome, whereas neither sequencing depth nor dataset had a significant effect (Table 2). The functional profiles of mountain gorilla microbiomes differed significantly from the other two subspecies (Table 2).

**Table 2.**
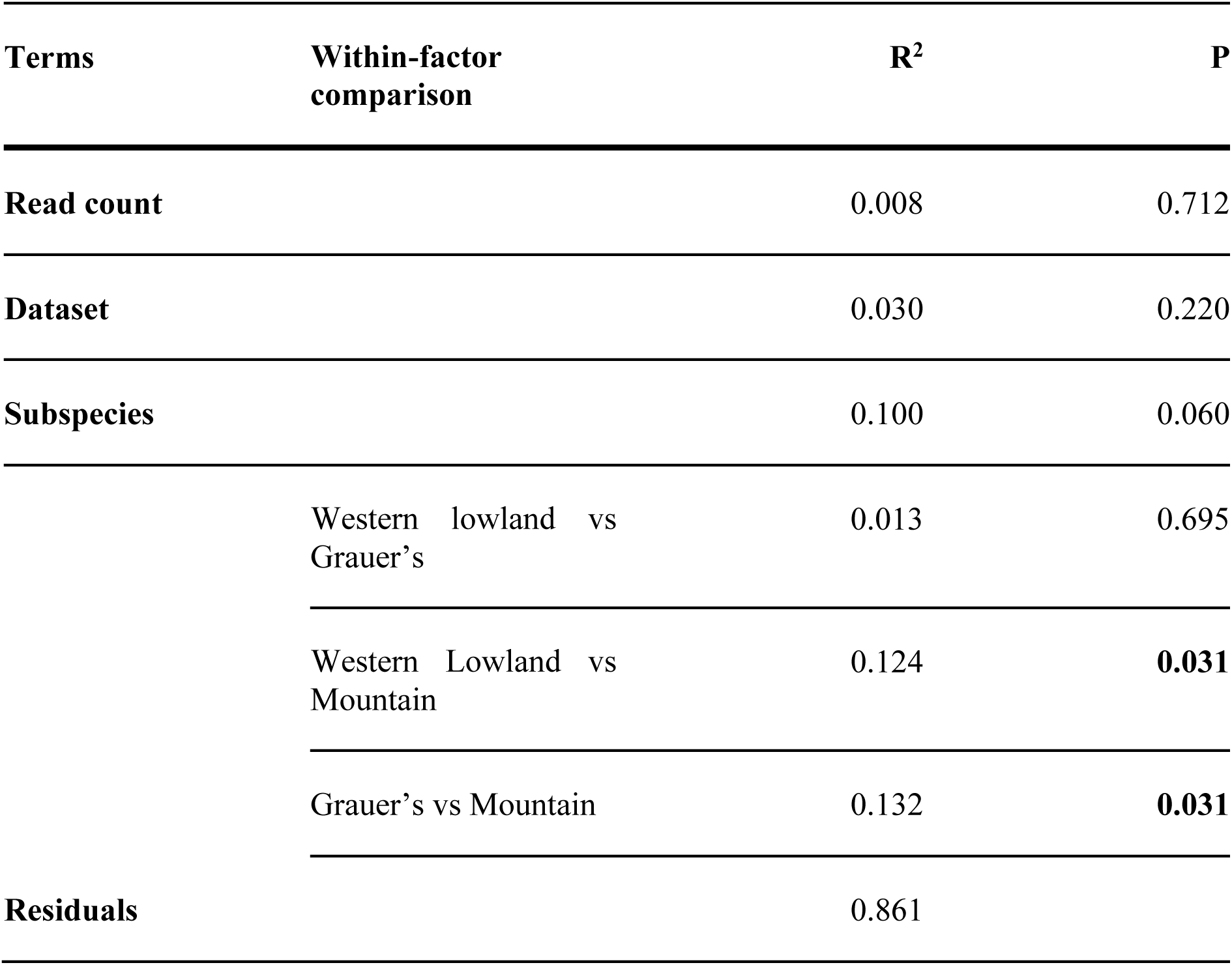
PERMANOVA results, showing the effect size (R^2^) and p-value of factors putatively contributing to the variance in abundance of biological processes among sampled microbiota. Pairwise PERMANOVA between subspecies show the effect size (R^2^) and p-values adjusted for false discovery rate. P-values below 0.05 are shown in bold.

| Terms | Within-factor comparison | $R^2$ | P |
| --- | --- | --- | --- |
| Read count |  | 0.008 | 0.712 |
| Dataset |  | 0.030 | 0.220 |
| Subspecies |  | 0.100 | 0.060 |
|  | Western lowland vs Grauer's | 0.013 | 0.695 |
|  | Western Lowland vs Mountain | 0.124 | **0.031** |
|  | Grauer's vs Mountain | 0.132 | **0.031** |
| Residuals |  | 0.861 |  |

We used ANCOM to identify differentially abundant biological processes in the dental calculus microbial communities of the gorilla subspecies, requiring that these processes were represented in at least 30% of the samples in at least one subspecies. Mountain gorillas stood out again, as they missed genes for 236 of the 262 differentially abundant processes (Table S6). In contrast, western lowland and Grauer’s gorillas were missing only 13 and 6 processes, respectively. Among six processes that were found in all three gorilla subspecies, mountain gorillas differed significantly from the others in four processes.

### Metagenome-Assembled Genomes

We applied an iterative assembly and binning approach (Methods) to resolve initial MAGs using decontaminated sequencing reads from all samples and museum controls. Across all samples, we constructed a total of eight high-quality (>90% completion, <5% contamination) and 27 medium quality MAGs (>50% completion, <10% contamination), belonging to 24 distinct bacterial families (Table S7). Among these 35 MAGs, 25 could be classified to the genus level, with two MAGs assigned to provisional/uncultured genera (UBA8133 and RUG013; Table S7). To place the MAGs within a broader evolutionary context, we constructed a phylogenetic tree including the MAGs and phylogenetically closest bacterial genomes from the Genome Taxonomy Database (GTDB; (Parks et al., 2018); Figure S4). However, for several MAGs assembled in this study we were unable to identify a closely related published genome.

Fourteen MAGs reconstructed from dental calculus samples were also present in museum controls (a gorilla skin sample, a gorilla petrous bone, a skull swab and a museum shelf swab). Eight MAGs were present only in the skin sample and only at low abundances (Figure S5). However, five MAGs were found at higher abundance in at least one of the museum controls compared to any gorilla dental calculus sample (Figure S5). This includes MAGs of *Erysipelothrix* (found in gorilla petrous bone), two different *Planococcaceae* (found in the petrous bone and museum shelf swab), as well as *Propionibacterium* and *Exiguobacterium* (both isolated from museum skull swab). These MAGs belong to genera that are common contaminants in metagenomic studies (i.e. *Propionibacterium*, which is a major genus of common skin bacteria (Salter et al., 2014)) and persistent in environmental reservoirs (i.e. *Erysipelothrix*; (Eriksson et al., 2014)). These five MAGs were excluded from downstream analyses. To further authenticate our reconstructed MAGs, we used a collection of isolation source records from public databases of bacterial ecological metadata (Table S8). Taxa with a high proportion (>=25%) of isolation records from sources categorised as environmental/contaminant were considered as potential contaminants (Table S8). Using this approach, we identified 10 additional MAGs that represent likely contaminants (Figures S5& S6).

The remaining 20 MAGs recovered here are members of the oral microbiome and are present in databases of common oral taxa (Chen et al., 2010; Fellows Yates et al., 2021). This includes taxa commonly associated with dental plaque communities: *Rothia* (Tsuzukibashi et al., 2017), *Olsenella* (Abusleme et al., 2013; Socransky et al., 1998), *Corynebacterium* (Mark Welch et al., 2016), *Lautropia* (Gerner-Smidt et al., 1994), *Neisseria (Donati et al., 2016)* and *Actinomyces* (Kolenbrander et al., 2010). *Rothia* species are particularly abundant in gorillas compared to humans and other non-human primates (Fellows Yates et al., 2021). Three other MAGs were present in at least 65% of all samples: a MAG most closely related to *Neisseria* (in 95.7% of samples) was found in all Grauer’s and mountain gorillas, and the MAGs characterised to the family *Actinomycetaceae* and the genus *Lautropia*, both of which were found in all samples of mountain gorillas (Figure S5).

For two MAGs of high completeness and high prevalence in our samples (*Neisseria* and *Limosilactobacillus gorillae*), we produced an alignment of core genes to further investigate how they relate to the known diversity of these taxa. The *Neisseria* MAG clustered with a subset of *Neisseria* species isolated exclusively from humans and was more divergent from *Neisseria* species that formed a clade with isolates from other animals (Figure 4a). Its phylogenetic placement suggests a distinct and not yet identified *Neisseria* taxon. We also recovered a near-complete MAG of *Limosilactobacillus gorillae* (Figure S7), a species previously isolated from faeces of mountain and western lowland gorillas (Tsuchida et al., 2018, 2015). The dental calculus *L. gorillae* MAG was sister to the gorilla faecal isolate but distinct from faecal isolates of other primates (Figure S7a).

**Figure 4:**
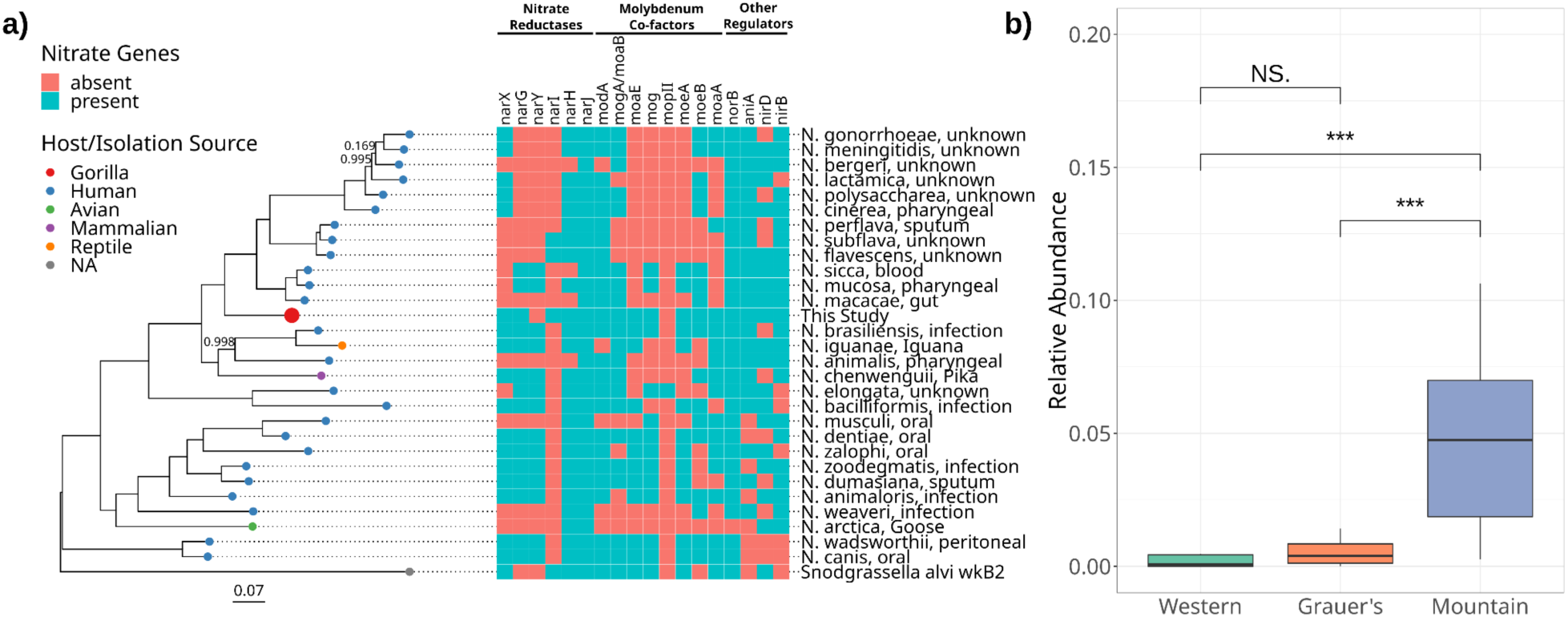
a) Maximum-likelihood phylogeny based on the alignment of core gene sequences from the *Neisseria* MAG and additional published genomes of the same genus. Tip points are coloured by host species identity, and tip labels include the isolation source for each published genome. Scale bar units are the number of substitutions per site and node values represent the proportion of node support out of 100 bootstrap replicates, with unlabeled nodes having complete support (1.0). To the right, the presence of key genes involved in the reduction of nitrate in the oral cavity is shown for each *Neisseria* species. **b)** Relative abundances (CLR transformed) of the *Neisseria* MAG in different gorilla subspecies. The results of a Wilcoxon test are denoted by brackets above boxplots (NS = not significant; *** = p > 0.001).

Oral bacteria play an important role in the metabolism of inorganic nitrate and its reduction to nitrite and nitric oxide, which is essential for the regulation of cardiovascular, metabolic, and neurological processes (Bryan et al., 2017; Hezel and Weitzberg, 2015). Several oral nitrate reducing bacteria, including *Rothia*, *Neisseria*, *Veillonella*, *Corynebacterium*, *Actinomyces*, *Selenomonas*, *Propionibacterium*, *Fusobacterium*, and *Eikenella (Rosier et al., 2022),* were present in gorilla dental calculus samples (Figure S5, Table S5). We used MAGs of *Rothia*, *Neisseria*, and *Veillonella*, which were prevalent in the study samples (Figure S5) to analyse the presence of nitrate-reducing genes in these members of the gorilla oral microbiome. Although we recovered a medium quality MAG of *Corynebacterium* from some samples of western and Grauer’s gorillas (Figure S5), it was less prevalent and abundant in gorilla dental calculus and is lacking well-characterised genes involved in nitrate reduction. For *Neisseria*, we could confirm the presence of five nitrate reducates (*nar*), seven genes that provide the cofactor molybdenum to the nitrate reductase enzymes, and four additional genes involved in the regulation of nitrate and nitrite reduction (*aniA*, *nirB*, *nirD*, *norB*; Figure 4a). Only two genes involved in nitrate reduction, *narY* and *mopII*, were not found in the gorilla dental calculus MAG. However, these genes are also absent from other nitrate-reducing oral bacteria (Rosier et al., 2020). Genomic data thus suggests that *Neisseria* isolated from gorilla dental calculus likely has the full capacity to reduce nitrate. The *Neisseria* MAG recovered in this study was significantly more abundant in mountain gorillas compared to the other host subspecies (Figure 4b).

*Rothia* and *Veillonella* possessed a smaller repertoire of nitrate-reducing genes compared to *Neisseria*. The *Rothia* MAG recovered in this study had several nitrate reductases and molybdenum cofactor genes, but they differed from genes present in *R. dentocariosa*, a bacterium well studied for its nitrate-reducing capacity (Figure S8; (Rosier et al., 2020). We were also able to identify nine genes involved in nitrate metabolism in *Veillonella*, a well known reducer of nitrate in the oral cavity (Hyde et al., 2014a) (Figure S8). Similar to *Neisseria*, *Veillonella* was significantly more abundant in mountain gorillas than in the other two gorilla subspecies, whereas the abundance of *Rothia* did not differ among gorilla subspecies (Figure S8b and d).

### Altitude may drive oral microbiome composition

Both taxonomic and functional analyses suggest that Grauer’s gorilla oral microbiomes are more similar to western lowland gorillas than to mountain gorillas, despite sharing a close evolutionary relationship and an adjacent geographic range with the latter. However, the distribution ranges of all three gorilla subspecies differ in elevation, with mountain gorillas occurring at the highest altitudes. As altitude can influence temperature, humidity and food diversity, which in turn can influence microbial communities, we performed partial Mantel tests between altitudinal distances and taxonomic (Jaccard/Aitchison distances) or functional (Euclidean distance) composition of the oral microbiome, while accounting for log-transformed geographical distance. Geographic location and altitude of the specimens were approximated based on museum records. However, when considering the entire dataset, we did not detect a correlation between altitude and either taxonomy or function (Mantel test: R ranging from -0.13 to 0.06, p>0.12).

Since Grauer’s gorillas occupy the widest altitudinal range of all gorilla subspecies, we divided Grauer’s gorilla samples into high- (>1000 masl, n=8) and low-altitude (≤1000 masl, n=8) groups and tested for differentiation of the oral microbiome both between these groups and among host subspecies. While no compositional differences were identified between the oral microbiomes of western lowland and low altitude Grauer’s gorillas using relative abundance measures, western lowland and high altitude Grauer’s gorillas displayed marginally significant differences based on pairwise PERMANOVA (FDR-adjusted p-value=0.075, Table 3). The oral microbiome of mountain gorillas consistently differed from other subspecies, independent of altitude grouping (Table 3). Yet, high-altitude Grauer’s gorillas showed considerable overlap with mountain gorillas in the first two axes of a PCoA based on Aitchison distances (Figure S9).

**Table 3.** Results of the pairwise PERMANOVA showing the effect size (R^2^) of comparisons of taxonomic composition based on Aitchison distances among four gorilla groups (western lowland, low-latitude Grauer’s [≤1000 masl], high-altitude Grauer’s [>1000 masl], and mountain gorillas) and the associated p-values (adjusted for false discovery rate). P-values below 0.05 are shown in bold.

| Comparison | $R^2$ | p-value (FDR-adjusted) |
| --- | --- | --- |
| Western Lowland vs Low-altitude Grauer's | 0.048 | 0.515 |
| Western Lowland vs High-altitude Grauer's | 0.070 | 0.075 |
| Western Lowland vs Mountain | 0.083 | **0.002** |
| Low-altitude Grauer's vs High-altitude Grauer's | 0.084 | 0.164 |
| Low-altitude Grauer's vs Mountain | 0.098 | **0.002** |
| High-altitude Grauer's vs Mountain | 0.087 | **0.002** |

To further investigate the effect of altitude on the oral microbiome, we identified a set of differentially abundant taxa between mountain and western lowland gorillas (n=41) and asked if Grauer’s gorillas from different altitudes show similarities with these two subspecies. We observe no significant difference between low-altitude Grauer’s and western lowland gorilla and between high-altitude Grauer’s and mountain gorillas, whereas all other comparisons were significant (p<0.003, Table S9).

### Dental calculus reflects dietary differences of gorilla subspecies

After removing reads assigned to bacterial and viral taxa, we performed taxonomic classification of the eukaryotic diversity present within our samples to characterise gorilla dietary components using Kraken2 with the NCBI ‘nt’ database. Despite applying extensive decontamination procedures (Methods) and removing taxa with few supporting reads (<10), we still recovered erroneous assignments, such as mollusks or other mammals, which gorillas are unlikely to consume (Mann et al., 2020). We therefore restricted our analyses to eukaryotic families that are known to be part of the gorilla diet ((Michel et al., 2022; Remis et al., 2001; Rogers et al., 2004; Rothman et al., 2014; Yamagiwa et al., 2005); Methods). The majority of the 371 genera (65 families) detected in our dataset (Figure S10) were plants (n=360), but we also identified insects, specifically ants, (N=7) and lichen-forming fungi of the *Parmeliaceae* family (N=6).

We detected 22 genera that differed in abundance across gorilla subspecies, all belonging to plants, except for the fungal genus *Parmotrema* (Figure 5, Table S10). The broad dietary patterns observed here agree with literature reports. For instance, bamboo - represented here by the African genus *Oldeania*, as well as Asian genera *Phyllostachys*, *Bambusa,* and *Ferrocalamus*, which are likely misclassifications of African bamboo species (Brealey et al., 2020) - was mainly detected in mountain and Grauer’s gorillas, as expected (Rothman et al., 2014; Yamagiwa et al., 2005); Figure 5). *Marantochloa* and *Thaumatococcus* of the *Marantaceae* family were detected in western lowland gorillas, which are known to consume them (Rogers et al., 2004), but also in Grauer’s gorillas, where the family *Marantaceae* has been reported as part of the diet (Michel et al., 2022). The family Fabaceae is consumed by all gorilla subspecies and the genus *Tamarindus* detected here is likely a misidentification of another member of this family, as *Tamarindus* itself has not been reported to be part of gorilla diet. Grauer’s gorillas appear to have the most diverse diet of the three subspecies, consuming foodstuffs typical of both western lowland and mountain gorillas (Figure 5). This is in accordance with the literature (Michel et al., 2022; Remis et al., 2001; Rogers et al., 2004; Rothman et al., 2014; Yamagiwa et al., 2005), which reports 61 plant families as being part of their diet, as opposed to 41 and 45 families for western lowland and mountain gorillas, respectively.

**Figure 5.**
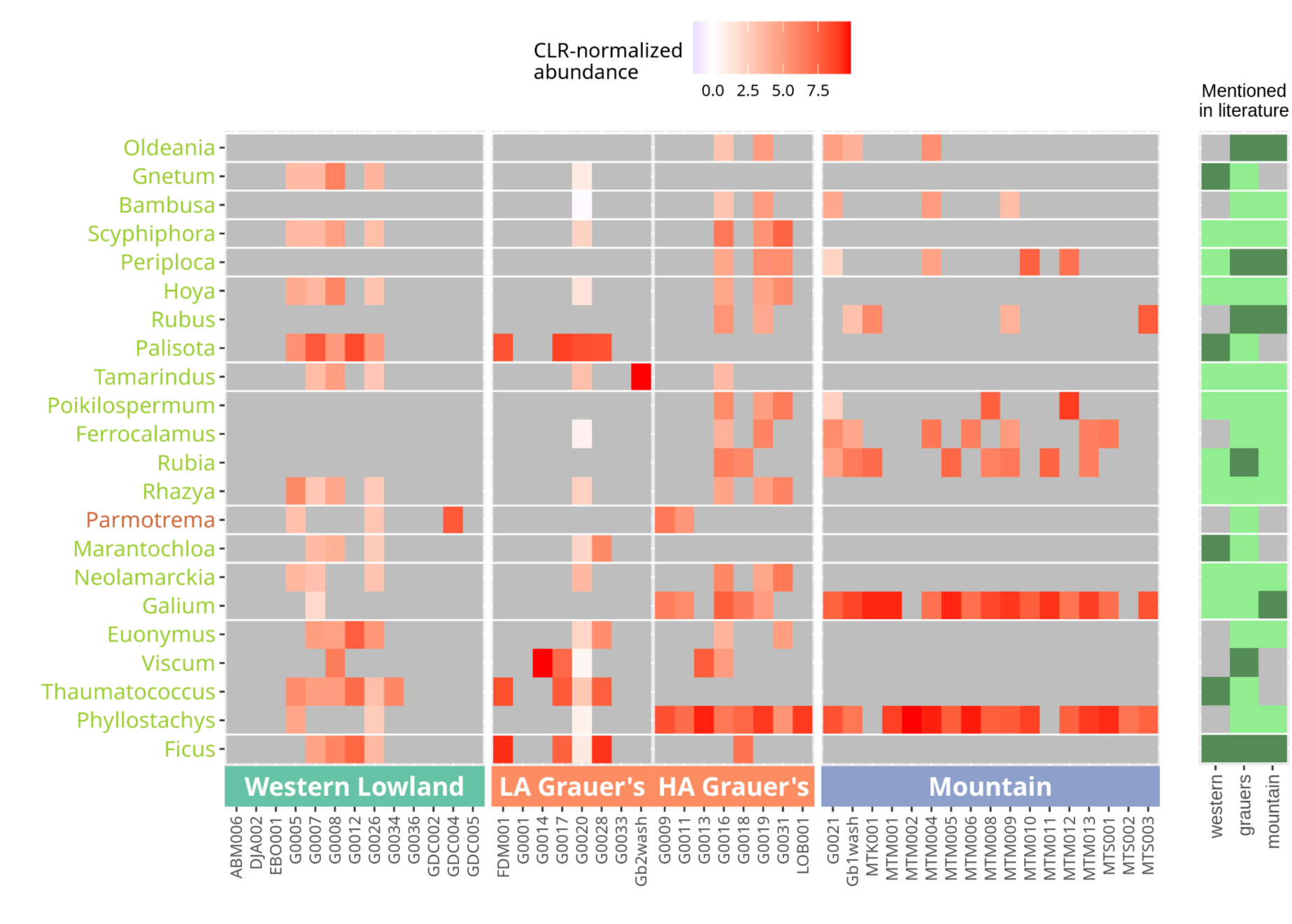
Heatmap based on normalised abundances for differentially abundant dietary taxa at the genus level. For clarity, taxa with abundance equal to 0 are shown in grey. The bars on the right indicate if the family (light green) or the specific genus (dark green) have been reported as part of the diet of each gorilla subspecies (grey: no mention). Horizontal white lines separate plant families.

Similar to our analyses of differentially abundant bacteria, dietary components which were differentially abundant between mountain and western lowland gorillas showed no significant difference between low-altitude Grauer’s and western lowland gorilla and between high-altitude Grauer’s and mountain gorillas, while significantly differentiating low- and high-altitude Grauer’s gorillas (Table S9).

## Discussion

Using three gorilla subspecies, we investigated how evolutionary relationships and ecological factors shape the dental calculus oral microbiome in closely related host taxa. Ecologically, all three subspecies differ from each other, but Grauer’s and mountain gorillas are geographically proximate and more closely related to each other than to western lowland gorillas. One of the strongest differences between the three subspecies is the altitude of their ranges, which in turn influences diet, reflected, e.g., in the level of frugivory. Western lowland gorillas occupy the lowest altitudes and are the most frugivorous (Rogers et al., 2004; Takenoshita and Yamagiwa, 2008). Mountain gorillas occupy the highest altitude and are the least frugivorous (Ganas et al., 2004; Maisels et al., 2018; Plumptre et al., 2016), whereas Grauer’s gorillas live across the largest altitudinal range and hence straddle the ecological conditions faced by both western lowland and mountain gorillas (Yamagiwa et al., 2008). Ecological conditions and dietary differences are expected to affect the host-associated microbial communities (Janiak et al., 2021; Kartzinel et al., 2019; Li et al., 2013; Wade, 2013; Youngblut et al., 2019).

By investigating the oral microbiome preserved in dental calculus, we show that oral microbial communities among closely related gorilla subspecies differ, both taxonomically and functionally. In particular, multiple lines of evidence show that the oral microbiome of mountain gorillas differs from the other gorilla subspecies in taxonomic composition, relative abundance, and functional profiles of microbial taxa (Tables 1 and 2), despite the close evolutionary relationship and geographic proximity between mountain and Grauer’s gorillas. The oral microbiome of Grauer’s gorillas shows strong similarity to western lowland gorillas, although the two species are more distantly related and separated by the Congo Basin. When considering differentially abundant microbial taxa and dietary profiles, Grauer’s gorillas appear intermediate, displaying a combination of taxa found in both western lowland and mountain gorillas (Figure 3 and 6). This suggests that host phylogenetic relationships are less important for the structure of the oral microbiome in closely related species than ecological conditions, possibly including diet.

These observations deviate from previous studies that identified a phylogenetic signal in the oral microbiome in wild animals. However, these studies compared distantly related species like chimpanzees, humans (Li et al., 2013), other great apes (Boehlke et al., 2020), and even more diverse groups of mammals (Brealey et al., 2020; Ozga and Ottoni, 2021). Often the effects of host phylogenetic relationships and ecology could not be decoupled. For instance, Smith et al. (2021) found differences among three snake species that each occupied a distinct ecological niche. Similarly, Soares-Castro et al. (2019) studied three marine mammals, among which the most evolutionary divergent species also lived in a different habitat, so the effects of ecology were indistinguishable from those of evolutionary relationships.

Yet, ecological effects on a short evolutionary scale, as observed in this study, do not preclude co-diversification of the host and its associated microbiome over longer evolutionary time. They rather point to the presence of rapid and dynamic responses of the host-associated microbial communities to environmental conditions, potentially through the seeding of the dental calculus oral microbiome at least to some extent with environmental taxa (see below). In addition, our results are in accordance with other studies that propose external factors, such as diet, influencing the oral microbiome (Adler et al., 2016; Hyde et al., 2014b; Janiak et al., 2021), including Li et al. (2013) who observed that the oral microbiome of captive apes is distinct from their wild counterparts and more similar to other captive apes, highlighting the importance of environmental factors/diet.

### Oral microbiome diversity might reflect ecology or diet

Our analyses provide a tentative suggestion that ecological differences, approximated here by altitude, may be the primary factor shaping oral microbiome diversity. Specifically, grouping Grauer’s gorillas into low- and high-altitude populations, we showed that low-altitude populations harbour an oral microbiome that is indistinguishable from western lowland gorillas, whereas high-altitude populations show near-significant differences (Table 3). Although the oral microbiome of Grauer’s gorillas is significantly different from mountain gorillas independent of the altitude, this may be because most mountain gorillas in our dataset were collected above 3000m, notably higher than any of the Grauer’s gorilla samples (600-2100m; Table S1). We also observe compositional differences between high- and low-altitude groups of Grauer’s gorillas when considering oral and dietary taxa that differentiate mountain and western lowland gorillas (Table S9). The high-altitude populations of Grauer’s gorillas appear indistinguishable from mountain gorillas, whereas low-altitude Grauer’s gorillas show no differences to western lowland gorillas, supporting the effect of ecological differences connected to altitude that shape microbial and dietary composition. Generally, the dietary profiles of Grauer’s gorillas appear to contain dietary items present in both western lowland and mountain gorillas (Figures 6 and S10). Our observation from dental calculus is in line with previous studies (Table S11), which report considerable overlap in the diet of Grauer’s gorillas with western lowland (30 of the 41 plant families) and mountain (30 of the 45 plant families) gorillas. In contrast, only 16 dietary families are consumed by both western lowland and mountain gorillas.

It is also intriguing to speculate that the local environment itself could affect the composition of the dental calculus community by serving as a source of at least some colonising taxa, for example via the consumed phyllosphere and rhizosphere. A recent study of the human oral microbiome found that specific taxa within the dental plaque community are evolutionarily close to environmental bacteria, whereas those living on the tongue surface are more closely related to other host-associated taxa (Shaiber et al., 2020). This finding suggested that dental plaque microbial communities could be more affected by the environment than other host-associated microbiomes. Within the gorilla oral microbiome, we detected microbial taxa that likely have a dietary origin and differ in abundance among host subspecies, possibly reflecting dietary differences. In particular, bacteria associated with the *Fabaceae* rhizosphere, including *Agrobacterium deltaense, A. fabacearum* (Delamuta et al., 2020; Yan et al., 2017), and three *Rhizobium* species (Poole et al., 2018) may be derived from the consumed dietary items or accidentally ingested soil. These bacteria were significantly more abundant in the dental calculus of western lowland and Grauer’s gorillas than in mountain gorillas (Figure 3, Table S5). Two lines of evidence provide a link between these microbial taxa and dietary differences among the gorilla subspecies. First, our data suggest a higher abundance of the family *Fabaceae* in western lowland and Grauer’s gorillas (Figures 6 and S10, Table S10) compared to mountain gorillas. Second, although no comparative data exists on the prevalence of roots and rhizomes in the diet of different gorilla species, indirect evidence suggests that western lowland gorillas consume more roots than mountain gorillas. Behavioural observations report frequent root and rhizome consumption in western lowland gorillas of all ages (Fletcher and Nowell, 2008) and comparative dentition analyses show increased tooth wear in western lowland gorillas compared to mountain gorillas, in which tooth wear is well explained by the time spent feeding on roots (Galbany et al., 2016).

Although dental calculus is a highly suitable material for the study of the oral microbiome, several previous studies have identified caveats associated with dietary analyses from this source material (Haas et al., 2022; Mann et al., 2020; Ottoni et al., 2019). Sample contamination, the low proportion of eukaryotic reads, and the sparsity of eukaryotic reference databases lead to biases and present a considerable challenge to the analysis of dental calculus. In addition, reference genome contamination can result in false positive taxonomic assignments (Mann et al., 2020). In the present study, the authenticity of putatively dietary taxa could not be assessed using damage patterns due to low abundance of eukaryotic reads. Nevertheless, we were able to detect dietary differences among gorilla subspecies, which are mostly in agreement with previous knowledge of gorilla diets (Figure 5) (Michel et al., 2022; Remis et al., 2001; Rogers et al., 2004; Rothman et al., 2014; Yamagiwa et al., 2005).

### Metagenome-assembled genomes add to the understanding of wild microbiomes

The recovery of high quality MAGs from metagenomic data can provide an important perspective on the evolutionary relationships among microbial taxa and between microorganisms and their hosts. As reference databases are often biased towards organisms of medical, agricultural, and industrial importance (Marcelino et al., 2020; Velsko et al., 2018), MAGs recovered from novel environments, such as dental calculus oral microbiome of non-human animals, may represent poorly described, understudied or completely unknown microbial lineages (Jiao et al., 2021). Several of the high and medium quality MAGs constructed in this study were recovered in higher abundance from environmental controls than from any dental calculus samples and were considered to represent environmental contaminants. However, other MAGs were highly abundant in the dental calculus samples and belong to taxa that are members of the oral microbiome (Figures S5 & S6). Many of these MAGs were distinct from the evolutionary closest sequenced genomes available in GTDB (Figures S4, S7, S8, Figure 4), suggesting that they may represent undescribed bacterial lineages, in line with discoveries of novel microbial diversity in unstudied host species (Levin et al., 2021; Youngblut et al., 2020). The closest evolutionary relatives of several bacterial taxa with high quality MAGs were found among isolates from primates, including humans (Figures 5a, S7, S8).

In this study, we recovered a near-complete MAG of *Limosilactobacillus gorillae*, which was more closely related to a faecal isolate from a captive gorilla than to a faecal isolate from another primate (Figure S7a). Usually associated with gorilla faeces (Tsuchida et al., 2014), the presence of *L. gorillae* within the oral microbiome may be the result of coprophagy, a common behaviour among wild gorillas (Graczyk and Cranfield, 2003). However, questions remain regarding the persistence of faecal-associated bacteria within dental calculus and their potential function within the plaque biofilm. Coprophagic behaviour is suggested to serve as a route for vertical or horizontal transmission of gastrointestinal microbiota between individuals (Abusleme et al., 2020) and may have a stabilising impact on host-associated microbial communities overall (Bo et al., 2020).

We also reconstructed MAGs of *Neisseria*, *Rothia*, and *Veillonella* and confirmed the presence of nitrate-reducing genes in these members of the gorilla oral microbiome (Figures 5a, S8). The reduction of nitrate to nitrite from dietary sources by oral bacteria provides the precursor to nitric oxide (NO), which is an important signalling and effector molecule (Jones et al., 2021). Along with multiple benefits to circulatory health (Lundberg et al., 2018), nitric oxide is increased in response to hypoxic stress (Feelisch, 2018; Levett et al., 2011), and results in increased blood oxygen levels in the host (Beall et al., 2012). Human populations adapted to high altitude exhale higher concentrations of nitric oxide compared to low-land populations (Beall et al., 2001; Erzurum et al., 2007). For these reasons, regulation of nitric oxide metabolism is thought to be beneficial to high-altitude adaptation (Beall et al., 2012).

Mountain gorillas live at elevation of up to 3,800 masl (Williamson et al., 2013) and hence are exposed to hypoxic stress (Ruff et al., 2022). They also show higher abundance of *Neisseria* and *Veillonella* compared to the other two gorilla subspecies, which live at considerably lower altitudes (Figures 5b and S8d). Although it is not known if dietary nitrate amounts differ among gorilla subspecies, the primarily herbivorous diet of mountain gorillas is naturally rich in nitrate. Taken together, it is likely that high abundance of nitrate-reducing oral taxa may aid mountain gorillas cope with the physiological demands of their high-altitude habitat. This finding is particularly intriguing, as comparative genomic analyses failed to uncover host-encoded genes related to high-altitude adaptation in mountain gorillas (Xue et al., 2015). Our results suggest an important role for host-associated microbiomes in promoting host adaptations.

### Mitochondrial DNA in dental calculus can aid taxonomic assignments of the host

The mitochondrial sequences recovered from the dental calculus metagenomes were used to confirm gorilla subspecies identity. Six of the samples yielded mitochondrial genomes with completeness >80% and coverage >3X (maximum 94% completeness and 77X coverage, Table S3). However, in many cases even a few mitochondrial fragments were sufficient to allow subspecies assignment. We could genetically distinguish gorilla species (eastern versus western gorillas) with mitochondrial genome coverage as low as 2.6% and eastern gorilla subspecies (mountain versus Grauer’s) with coverage as low as 6.6%. Grauer’s and mountain gorilla mitochondrial genomes could be successfully distinguished despite differing by only 0.5% (72 positions, excluding the hypervariable D-loop region (van der Valk et al., 2018)). This observation supports the usefulness of dental calculus as material for obtaining genetic information about the host (Mann et al., 2018; Ozga et al., 2016; Warinner, 2016; Warinner et al., 2015).

## Conclusions

Our study is, to our knowledge, the first to investigate the evolution of the dental calculus oral microbiome at the early stages of the speciation process of the host and adds to the new but growing field of research on the microbiomes of wild animals. We find that in closely related species, evolutionary relationships are less important than ecology in explaining taxonomic composition and function of the oral microbiome. Host-associated microbial communities have been proposed to contribute to host adaptation, partly because they can respond rapidly to changing environmental conditions (Alberdi et al., 2016). We find that taxa enriched in the mountain gorilla oral microbiome may facilitate their high-altitude lifestyle through increased nitrate reduction potential and the associated physiological benefits. Our discovery of distinct phylo- and rhizosphere taxa in the dental calculus of different gorilla subspecies suggests a close connection between this host-associated microbiome and the local environment. Colonisation of the host by environmental taxa with beneficial functions in local adaptation and health may present an exciting evolutionary route to be explored in the future.

## Methods

### Sample collection

The study dataset consisted of dental calculus samples from 57 specimens of three gorilla subspecies: 16 western lowlands gorillas, 22 Grauer’s gorillas, and 19 mountain gorillas (Table S1). Dental calculus samples of western lowland gorillas were collected at the Royal Museum for Central Africa (RMCA, Tervuren, Belgium) and the Cleveland Museum of Natural History (CMNH, USA). Samples of mountain gorillas came from the Swedish Museum of Natural History (Naturhistoriska Riksmuseet - NRM, Stockholm, Sweden) and the RMCA. Samples of Grauer’s gorillas were collected primarily at RMCA, with a few samples from NRM and the Royal Belgian Institute of Natural Sciences (RBINS, Brussels, Belgium). Our dataset consisted of newly generated shotgun data from 26 specimens and published gorilla dental calculus sequences: two samples from Brealey et al. (2020) and 29 samples from Fellows Yates et al. (2021). We also included sequences from 35 published and newly generated extraction blanks, twelve library preparation blanks, and four museum controls. One extraction and one library blank had too few reads for taxonomic classification and were therefore excluded from downstream analysis (Table S1). The museum controls consisted of two swabs taken at NRM from a museum shelf surface holding reindeer (*Rangifer tarandus*) specimens and from the surface of a brown bear (*Ursus arctos*) skull. In addition, we included data from two gorilla specimens from previous studies (van der Valk et al., 2019, 2017): A petrous bone sample from a specimen at NRM and a skin sample from RMCA (ENA accession numbers: ERR2503700 and ERR2868193, respectively). These sequences were generated to study the gorilla host, however, we removed host reads and retained microbial reads for our analyses.

### Preparation of Genomic Libraries and Metagenomic Shotgun Sequencing

All samples were processed in cleanroom facilities following appropriate methods for working with ancient and historical DNA. DNA was extracted according to the protocol by Dabney et al. (2013) with slight changes, as described in Brealey et al. (2020) and Fellows Yates et al. (2021). The datasets differed in library preparation protocol, indexing strategy, sequencing platform, read length, and sequencing depth (Table S1). Samples from four specimens were processed in both laboratory facilities: Uppsala University, Uppsala, Sweden and the Max Planck Institute for the Science of Human History, Jena, Germany (Fellows Yates et al., 2021). We used these technical duplicates to assess putative batch effects (Fig. S3), but only retained one sample per pair for biological analyses, selecting the one with the largest number of reads.

For the newly generated data, we followed the protocol as detailed in Brealey et al. (2021). Briefly, dental calculus samples ranging in weight from < 5 mg and up to 20 mg were surface-decontaminated using UV light (10 min at 254 nm) and washing in 500 µl of 0.5M ethylenediaminetetraacetate (EDTA) for 1 min (Brealey et al., 2021, 2020; Ozga et al., 2016). DNA was extracted using a silica-based method (Dabney et al., 2013) in batches of at most 16 samples with two negative controls. DNA was eluted in 45 µl of EB buffer (10 mM tris-hydrochloride, pH 8.0; QIAGEN, Netherlands) supplemented with 0.05% (v/v) Tween-20.

Double-stranded genomic libraries for all newly generated and previously sequenced samples were prepared following the double indexing protocol (Dabney and Meyer, 2012; Meyer and Kircher, 2010). Newly generated libraries and those previously published by (Brealey et al., 2020) included double in-line barcodes to guard against index hopping (van der Valk et al., 2020). A detailed protocol for library preparation is provided in Brealey et al. (2020). Library blanks were included for each batch of approximately 20 samples. Adapter-ligated libraries were quantified by a quantitative PCR assay, allowing us to estimate an appropriate number of index PCR cycles, which ranged from 8 to 20. Following indexing, individual libraries were purified with Qiagen MinElute columns and quantified using a quantitative PCR assay. All extraction and library blanks consistently showed lower DNA content than calculus samples (Table S1). We pooled 1.5 µl of each indexed library (including blanks and museum controls) and performed size selection with AMPure XP beads (Beckman Coulter, IN, USA) for fragments of approximately 100-500 bp in length. The pooled library was sequenced by SciLifeLab Uppsala on two Illumina NovaSeq S2 flowcells using paired-end 100 bp read length and V1 sequencing chemistry.

### Preprocessing of sequencing data

For each sample, the data generated from different sequencing runs was concatenated into a single file (separately for reverse and forward reads) and poly-G tails, which are artefacts of the two-colour sequencing chemistry of the NextSeq and NovaSeq Illumina platforms, were removed using fastp (V0.20.0; Chen et al., 2018). We removed unpaired reads with BBTools ‘repair.sh’ (V38.61b; Bushnell, 2014). For newly generated data, we used AdapterRemoval (V2.2.2; Schubert et al., 2016) to clip adapters, trim reads based on minimum phred quality (>=30) and length (30bp), and merge forward and reverse reads. The unmerged reads were excluded from downstream analysis (Table S1). In-line barcodes in newly generated data were trimmed from both ends using a Python script (Brealey et al., 2020). Read quality filtering was performed using PrinSeq-Lite (V0.20.4; Schmieder and Edwards, 2011) with a mean base quality threshold of 30. PCR duplicates were removed using a Python script (Brealey et al., 2020), which randomly kept one read among those with the same sequence. All commands and scripts used in our bioinformatic analysis are available at 10.5281/zenodo.6861585.

Sequences of the *phi*X bacteriophage, which is used as an internal control for Illumina sequencing (Mukherjee et al., 2015), were removed from the dataset by mapping against the *phiX* genome (GenBank: GCA_000819615.1) using BWA-MEM (V0.7.17; Li and Durbin, 2009). Unmapped reads were retained using SAMtools (V1.12; Danecek et al., 2021) and converted to FASTQ for downstream analysis, using the ‘bamtofastq’ function of BEDtools (V2.29.2; Quinlan and Hall, 2010). To remove host reads and potential human contamination, we mapped all the reads to a combined file containing western lowland gorilla and human reference genomes (GenBank: GCF_000151905.2 and GCF_000001405.38, respectively), repeating the steps above. Extraction blanks, library blanks, and museum controls were only mapped to the human genome. The unmapped reads were used in oral microbiome and dietary analyses, whereas the mapped reads were extracted, sorted, and indexed using SAMtools, and used for the host genomic analyses.

### Host genome analysis

#### Mitochondrial genome analysis

From the host mapped BAM files, we extracted all reads mapping to the gorilla mitochondrial genome from position 1 to position 15,446, with a mapping quality of 30 or higher, using SAMtools. Nucleotide positions 15,447-16,364, which contain the D-loop, were excluded to prevent the hypervariable regions from introducing biases in our analyses. We calculated sample coverage and the total number of reads mapping to the mitochondrial genome with SAMtools and generated a consensus sequence in FASTA format with ANGSD (V0.933; (Korneliussen et al., 2014).

#### Subspecies assignment verification

The subspecies identity of each sample was initially assigned using museum records. We confirmed these assignments using diagnostic sites within the host mitochondrial genome. Using 102 published mitochondrial genomes (Das et al., 2014; Hallast et al., 2016; Hu and Gao, 2016; van der Valk et al., 2018; Xu and Arnason, 1996); Table S12), we identified two sets of diagnostic sites that were fixed for different alleles in each of the contrasted groups using a custom Python script. The first set was species-specific, identifying membership in western versus eastern gorillas. The second set was subspecies-specific, distinguishing between mountain and Grauer’s gorillas.

#### Molecular sexing of samples

We used sexassign (Gower et al., 2019) to determine the sex of each individual (Table S1). To this end, we mapped filtered reads, prior to removing host and human reads, to the gorilla reference genome (GCF_000151905.2) and counted reads mapping to the X chromosome and the autosomes. sexassign conducts a likelihood ratio test comparing the observed X-to-autosome ratio to expected ratios for males (RX ∼0.5) and females (0.8 > RX < 1.2).

### Initial taxonomic classification and decontamination

#### Taxonomic classification with Kraken2/Bracken

After removing host and human reads, the retained unmapped reads were taxonomically classified using Kraken2 (V2.1.1; Wood et al., 2019) with default parameters. We used the standard Kraken2 database (accessed by the software on 1st September 2021), which includes all bacterial, archaeal and viral genomes from NCBI. Abundances were re-estimated at the species level using Bracken (V2.6.2; Lu et al., 2017), by redistributing reads from higher taxonomic levels to the species level, (setting read length to 55 and the rest of parameters in default). The average read length was set to 55 bp, which corresponded to the average of the mean read length of our data. The raw species table and the corresponding metadata were combined into a phyloseq object (V1.34.0; (McMurdie and Holmes, 2013) in R (V4.0.4; R Core Team, 2015). Taxonomic information was assigned based on taxonomic IDs from the NCBI taxonomy database using the ‘classification’ function of the taxize package (V0.9.99; (Chamberlain and Szöcs, 2013).

#### Removal of low-quality samples and contaminant taxa

Samples containing less than 300,000 processed reads were excluded from downstream analysis (for the distribution of read counts per sample see Figure S11). We then evaluated the composition of the microbial communities of the retained samples with FEAST (V0.1.0; Shenhav et al., 2019) using default parameters.

FEAST makes use of user-provided reference microbiomes to partition the composition of the study samples and to estimate the proportional contribution of the reference “sources”. We provided 29 such sources sampled from six different environments that were chosen to reflect both the oral cavity and potential contaminants. Specifically, we included human calculus (n=5; Mann et al., 2018) and human plaque (n=5; Human Microbiome Project Consortium, 2012; Lloyd-Price et al., 2017) as oral sources, and human gut (n=5; Human Microbiome Project Consortium, 2012; Lloyd-Price et al., 2017), human skin (n=5; Oh et al., 2014), tundra soil (n=5; Johnston et al., 2016) and laboratory contaminants excluding human sequences (n=4; Salter et al., 2014) as potential contaminants (Table S13). Gorilla dental calculus samples for which the proportion of the metagenomic community similar to the human oral microbiome (human calculus and human plaque considered jointly) was lower than 3%, according to the output of FEAST, were excluded from further analyses. The threshold was decided based on the distribution of the oral proportions in the study samples (Figure S3b).

For the retained samples, we then carried out a multi-step approach to remove putative contaminant microbial taxa. First, we used the R package decontam (V1.10.0; Davis et al., 2018) with the ‘isContaminant’ function (method=“combined”, normalize = TRUE). The package employs two main approaches: a prevalence-based approach, which identifies contaminants based on their increased prevalence in blanks, and a frequency-based approach, which relies on the observation of a reverse relationship between taxon abundance and DNA quantity in the sample. We used a combination of both decontamination approaches, performed separately for the subset of newly generated data, which also included two previously published samples processed in the same facility (Uppsala University, Sweden; Brealey et al., 2020), and data from gorilla dental calculus published by Fellows Yates et al. (2021) and processed at the Max Planck Institute for the Science of Human History, Germany. We refer to these datasets as ‘newly generated’ and Fellows Yates et al. (2021), respectively. Thresholds were allowed to differ by dataset (0.2 for newly generated and 0.3 for Fellows Yates et al. (2021)) and selected to minimise the number of oral taxa (taxa reported as oral in Chen et al. (2010) and Fellows Yates et al. (2021)) among all taxa identified as contaminants (Figure S12). All taxa identified as contaminants were then removed from the full dataset, regardless of which subset they were identified in.

Second, we performed abundance filtering on the entire dataset, setting each taxon to 0 if it had relative abundance <0.005% in a given sample. We chose this threshold, as it retained the highest number and proportion of oral taxa, while excluding known contaminant taxa (Table S2). Third, we considered the relative abundances of taxa in the museum controls to remove likely environmental contaminants. A microbial taxon was removed as a likely contaminant if it was found to have a higher relative abundance in at least one of the four museum controls than in any of the samples. Finally, we considered bacterial genera identified as common contaminants in typical molecular and specialised ancient DNA laboratories (Salter et al., 2014; Weyrich et al., 2019). Some of the listed genera included taxa found in the Human Oral Microbiome Database (HOMD; accessed July 14 2021; (Chen et al., 2010) and in the core hominid oral microbiome (Fellows Yates et al., 2021). Therefore, we used a two-step approach to remove these potential contaminant taxa. We directly removed genera that were listed only as contaminants and did not appear in the oral databases. Taxa that were present in both contaminant lists and the oral databases and had a sufficient number of reads in our dataset were investigated further by running mapDamage2 (V2.0.9; Jónsson et al., 2013). To this end, we identified the sample with the highest abundance of a given taxon (according to Kraken2 output, requiring at least 10,000 reads) and mapped the reads to the respective reference genome. Taxa with too few reads to be tested were automatically retained. Since many taxa needed to be investigated in this way, making a visual inspection impractical, we assessed the presence of damage using a custom R script (Data and Materials). The presence of typical post-mortem DNA damages was assumed if deamination (C-to-T in the 3’ end and G-to-A in the 5’ end) was the most frequent change for at least two of the three terminal positions for each fragment end and had the frequency above 0.02 for at least one position. This rule was employed because in-line barcodes can create atypical DNA damage patterns in metagenomic data, showing lower damage at 1st terminal position (Brealey et al., 2020). Bacterial taxa with no damage were assumed to be modern contaminants and were therefore removed. The resulting dataset was used for downstream taxonomic analyses.

#### Sequence-level decontamination

We applied additional filtering by removing sequencing reads from taxa identified as contaminants (n=167) to allow for functional characterization of microbial communities and reconstructing metagenome-assembled genomes (MAGs). To this end, we used two approaches. First, we removed reads assigned to contaminants using the ‘extract_kraken_reads.py’ script from the KrakenTools suite (V1.2; Wood et al., 2019). Using the read-level taxonomic assignments by Kraken2, we removed reads assigned to the 167 contaminant taxa identified above. This approach retained sequencing reads of all taxa not identified as contaminants (i.e. those included in the final taxonomic dataset), but also a large number of unclassified reads.

The reads which remained unclassified by Kraken2 could contain reads belonging to contaminant taxa. Therefore, we performed a further decontamination step. We constructed a dataset containing the genomes of the identified contaminant taxa and a selection of abundant non-contaminant taxa from our dataset. To identify non-contaminant taxa, we selected all taxa with more than 10,000 reads in at least one sample (as long as they were not identified as contaminants), for a total of 72 unique taxa. The non-contaminant taxa were included to avoid forcing the reads to map to the contaminants. For both contaminants and non-contaminants, one genome per taxon was selected from the NCBI assembly summary files for bacteria, archaea, and viruses (NCBI ftp accessed 14 April 2021, available at https://github.com/markella-moraitou/Gorilla_dental_calculus/RD2_mapping/) with the following order of priority: genome representation (full > partial), RefSeq (yes > no), assembly level (full > chromosome > scaffold > contig), publication date (most recent). Twenty contaminant, and 11 non-contaminant taxa did not have a reference genome available in NCBI assembly. For them, we obtained a different genome from the same genus, unless that genus was already represented, using the taxize R package, and selected one genome per missing taxon as a replacement.

Finally, all references (consisting of 155 contaminant genomes and 61 non-contaminant genomes) were concatenated into a combined database. We mapped the FASTQ files from the previous decontamination step to this combined reference using BWA-MEM with default settings and then used SAMtools to retain the unmapped reads and reads that mapped primarily to the non-contaminant genomes.

### Taxonomic analyses

All taxonomic analyses were performed at the species level. To account for the compositional nature of the data, a centred-log ratio (CLR) normalisation was applied (Gloor et al., 2017) using the ‘transform’ function of the microbiome R package (V1.12.0; Lahti and Shetty, 2018).

#### Oral composition

To assess differences in oral microbiome composition by subspecies and sex, we used two distance measures: Jaccard distances (Jaccard, 1901), which only consider presence-absence of taxa, and Aitchison’s distances (Aitchison and Aitchison, 1986), which also take into account relative abundances and are calculated as Euclidean distances on the CLR-transformed data. We performed PERMANOVAs using these two distance metrics by applying the ‘adonis’ function of the vegan package (V2.5-7; Oksanen et al., 2011) and setting the parameter ‘method’ to ‘jaccard’ and ‘euclidean’ for the non-normalised and normalised data, respectively. Because of the unequal distribution of host subspecies among the two datasets (newly generated versus Fellows Yates et al. (2021); Figure S1), we included dataset as a factor in each model and considered it prior to evaluating the contribution of biological variables. We used the ‘adonis.pair’ function of the EcolUtils R package (V0.1; Salazar, 2019) to perform pairwise PERMANOVAs for significant factors using Jaccard and Aitchison distance matrices produced with the ‘vegdist’ function of the vegan package. Principal Coordinate Analysis (PCoA) plots were generated using the ‘plot_ordination’ function of the phyloseq package. Because sex assignment was missing for some samples (Table S1), we re-ran PERMANOVAs for each distance measure for the subset of samples that could be sexed.

#### Alpha diversity

We used the ‘estimate_richness’ function of the phyloseq package to estimate alpha diversity with two different metrics, applied to the non-normalized data: the Chao1 estimator (Chao et al., 2009), which evaluates species richness while accounting for the number of species that are missing from the dataset, and the Shannon index (Shannon, 1948), which reflects evenness. To assess the effect of the host subspecies on alpha diversity, we ran an ANOVA model and confirmed the model met assumptions of normality, using a Shapiro-Wilk’s test on the Studentized residuals, and visually assessed the fit against the residuals (Quinn and Keough, 2002). Both Chao1 and Shannon models were transformed to account for skew (‘sqrt(500-x)’ and ‘exp(x)’, respectively). Finally, we used a Tukey post-hoc test to compare the different levels of the factor(s) that were statistically significant.

#### Identification of differentially abundant taxa

We identified bacterial taxa that significantly differed in abundance between host subspecies using ANCOM-II (Mandal et al., 2015), a method appropriate for compositional data, which accounts for the sparsity of the taxon table (Kaul et al., 2017). To account for the association between host subspecies and dataset (newly generated versus Fellows Yates *et al*. (2021); Figure S1) and to reduce the possibility that we identify dataset-specific taxa as differentially abundant, we ran the analysis twice. First, we identified microbial taxa that differed in abundance between host subspecies, while accounting for read depth. Second, we identified microbial taxa that differed in abundance between datasets, while accounting for the host subspecies. Only genera that differed among subspecies and did not differ by dataset were considered relevant for evaluating microbiome differences among gorilla subspecies. Differentially abundant taxa were identified as those above the 0.9 quantile of the ANCOM W-statistic and Kruskal-Wallace post-hoc test (Bonferroni adjusted p-values < 0.05), or those that were structurally absent from one or more subspecies. Only microbial taxa identified at the genus or species level were considered.

Following this procedure, we also used ANCOM to identify taxa that differed in abundance between mountain and western lowland gorillas, for both microbial and dietary taxonomic datasets. These taxa were then used to explore the effects of altitude, separating Grauer’s gorillas into high- and low-altitude populations and performing comparisons using both Jaccard and Aitchison distances in pairwise PERMANOVAs, in which differences in dataset identity were included as factors.

### Functional analysis

Decontaminated metagenomic reads were functionally characterised using an assembly-free approach implemented in the HUMAnN2 pipeline (V0.11.2; Franzosa et al., 2018) using the MetaPhlAn2, UniRef50, and ChocoPhlAn databases (Segata et al., 2012; Truong et al., 2015). The output tables containing the abundances of gene families with specific function were regrouped to show gene ontology (GO) terms (Ashburner et al., 2000; Gene Ontology Consortium, 2021), normalised to copies per million reads and feature abundances across taxa were aggregated in a single feature table, which was imported into R for further analysis. We subset our dataset to GO terms referring to biological processes and used PERMANOVA on their Euclidean distances to investigate functional differences of the oral microbiome by host subspecies. We then compared the abundances of biological processes among host subspecies using ANCOM, using the same approach as above.

### Metagenome-assembled genomes (MAGs)

We used the metaWRAP pipeline (V1.3.2; Uritskiy et al., 2018) to construct MAGs from decontaminated sequencing reads. To increase the amount of data available for building the initial assemblies with MEGAHIT (Li et al., 2016), we concatenated the reads across all samples. The initial assembly was split into genomic bins using three different metagenomic binning tools (metabat2; V2.9.1 (Kang et al., 2019), maxbin2; V2 2.2.4 (Wu et al., 2016), CONCOCT; V0.4.0 (Alneberg et al., 2013)) and we retained only those bins that were consistently defined by all three tools. For each MAG, we assigned taxonomic identity with GTDB-TK (Chaumeil et al., 2019) and constructed a phylogeny containing our MAGs and closely-related reference genomes identified by GTDB-TK. The abundance of MAGs in each sample was estimated using salmon (V0.13.1; Patro et al., 2017) and the CLR-transformed abundances were visualised using pheatmap (Kolde and Others, 2012). For each assembled and taxonomically identified MAG, we extracted and curated records of isolation sources (sources listed in Table S8) following an approach described by Madin et al. (2020). This information was used to categorise the isolation sources for taxa into one of the following categories: “contaminant”, “environmental”, “host-associated”, “oral”, or “unknown”.

To evaluate evolutionary relationships for select MAGs, reference genomes or gene sequences for the identified species/genus were downloaded from GenBank and Genome Taxonomy Database (GTDB; V202; (Parks et al., 2018). We created alignments using the amino acid sequences of core genes with PhyloPhlAn (V.3.0.2; Segata et al., 2013) and constructed maximum-likelihood phylogenies using FastTree (V2.1.10; Price et al., 2010), which we visualised using ggtree (V3.1.0; Yu et al., 2017).

We used two different approaches to determine the presence of genes involved in the nitrate reduction pathway in MAGs belonging to oral genera with nitrate reduction ability (see references provided in Rosier et al. (2020)). First, we used the pangenomic tool Panaroo (Tonkin-Hill et al., 2020) to determine presence of genes across a set of fragmented MAG assemblies. Second, we performed a translated BLAST search (tblastn, V2.11.0; Camacho et al., 2009) using protein sequences obtained from UniProt against MAG contigs. Genes in MAGs and reference genomes were considered as present if detected with either method.

### Estimating ecological variation in oral microbiome composition

Using altitude as a proxy for ecology and diet, we estimated the correlation between altitudinal, geographical and microbiome distances. First, we approximated altitude and geographic location (latitude, and longitude) for each specimen based on the collection locality as retrieved from museum records. We constructed Euclidean distance matrices for altitudinal and geographical distance using R package vegan. Using the ade4 R package (1.7-16; Dray et al., 2007), we performed partial Mantel tests (10,000 permutations) between microbiome distance matrices (Pearson correlation used for Jaccard distances and Spearman’s rank coefficient correlation for Aitchison taxonomic and Euclidean functional distances) and altitudinal distances, while accounting for the effect of log-transformed geographical distances.

To further investigate the effects of altitude on Grauer’s gorilla microbiomes, which is the subspecies with the largest altitudinal range (Plumptre et al., 2016), we grouped the Grauer’s gorilla samples into low- (500-1000 m above sea level [masl]) and high-altitude (>1000 masl) populations. We evaluated the contribution of altitude by performing a PERMANOVA, after accounting for potential dataset and host subspecies effects. Post hoc tests were performed using pairwise PERMANOVA, as described above.

### Dietary Analysis

Eukaryotic reads in dental calculus were used to identify taxa potentially consumed by gorillas. Before taxonomically classifying these reads, we removed reads assigned to prokaryotes and viruses from the metagenomic dataset using the ‘extract_kraken_reads.py’ script from KrakenTools. The resulting FASTQ files were used as input in a second run of Kraken2 using the NCBI ‘nt’ database (accessed 27 September 2021). A joint feature table containing species and genus-level assignments was produced using kraken-biom and imported into R for further analysis. The genome coverage for these taxonomic classifications was insufficient for verification using mapDamage2.

#### Preprocessing of feature table

The feature table produced by kraken-biom was used in a phyloseq object along with a corresponding taxonomic matrix, created using the taxize R package. Using this matrix, we aggregated all identified taxa to the genus level, and used the decontam (V.1.10.0; Davis et al., 2018) R package to remove likely contaminants, as described above for the oral microbiome analyses, including using different cut-offs for each dataset. Then, all genera present in the museum controls or with fewer than 10 reads per sample were removed, to reduce contaminants, spurious and unreliable classifications. Despite this filtering, spurious classifications or contaminants were still present (species that are unlikely to be consumed by gorillas, e.g. mollusks), so we restricted it to genera belonging to taxonomic families that were previously reported to be consumed by gorillas (Michel et al., 2022; Remis et al., 2001; Rogers et al., 2004; Rothman et al., 2014; Yamagiwa et al., 2005). Since only few reference genomes of tropical plants are available and classically-used barcoding genes represent only a small part of the genome, accurate taxonomic classification from a sparse metagenomic sample is difficult and can likely only be achieved on genus or family level (Mann et al., 2020). For this reason, we focused our analysis not only on genera known to be consumed by gorillas, but also included all detected genera in the same family. The resulting feature table was used to detect differentially abundant dietary components with ANCOM using the same approach as described above, while accounting for differences in the number of reads per sample.

#### Collecting gorilla diet reference data

We produced a reference dataset of gorilla foods (Table S11) by searching the literature for plant species known to be consumed by the three gorilla subspecies based either on direct observations, analyses of food remains and faeces or more recently on molecular evidence (Michel et al., 2022; Remis et al., 2001; Rogers et al., 2004; Rothman et al., 2014; Yamagiwa et al., 2005). We retrieved the corresponding taxonomic IDs from NCBI using the taxize R package. For the taxa that had no match in the NCBI Taxonomy database, we manually searched the database and corrected the names that were obviously misspelt (difference of 1-2 characters from the NCBI entry). For the remaining taxa, we manually searched the GBIF species database (Wheeler, 2004) and obtained GBIF taxonomic IDs. However, several taxa (26 out of 314) could not be found in either database and therefore were excluded from our analysis. The taxonomic IDs obtained from either NCBI or GBIF were used with the ‘classification’ function of taxize to obtain the genus and family name for each taxon.

## Availability of Data and Materials

Sequencing data generated in this study have been deposited on ENA, under Project Accession Number: PRJEB49638. Scripts for both preprocessing and downstream analysis are available at 10.5281/zenodo.6861585.

## Authors’ contributions

JCB, MM, AF extracted and prepared genomic libraries for the newly generated data. MM processed sequencing data, performed bioinformatic and statistical analyses (microbiome taxonomic and functional analyses, dietary and host genome analyses, altitudinal analyses) and wrote the manuscript draft. AF performed bioinformatic and statistical analyses (metagenome-assembled genomes, functional analyses, altitudinal analyses), wrote the respective parts of the manuscript and contributed to others. JCB performed initial bioinformatics analyses on a smaller dataset. JAFY and CW provided genomic data. KG conceived the research plan, interpreted the results, provided financial support and infrastructure, and contributed significantly to manuscript writing. All authors contributed to writing the final version of this manuscript.

## Acknowledgments

We would like to thank Henrique Leitão, Thijs Hofstede, Franziska Aron, and Cody Parker, who contributed with processing the samples in the ancient DNA lab, Tom van der Valk, Axel Jensen and Samantha López Clinton for their contribution to subspecies verification using mitochondrial DNA. We thank Daniela C. Kalthoff at the Swedish Museum of Natural History, Emmanuel Gilissen at the Royal Museum for Central Africa, Tom Geerinckx and Patrick Semal at the Royal Belgian Institute of Natural Sciences, and Lyman Jellema at the Cleveland Museum of Natural History for providing access to and assistance with the gorilla specimens. Sequencing was performed by the SNP&SEQ Technology Platform in Uppsala. The SNP&SEQ Technology Platform is part of the National Genomics Infrastructure Sweden and Science of Life Laboratory. The SNP&SEQ Platform is also supported by the Swedish Research Council and the Knut and Alice Wallenberg Foundation. We also acknowledge the National Bioinformatics Infrastructure for providing computational resources to this project.

