## Supplemental Figures S1-S12 for "Ecology, not host phylogeny, shapes the oral microbiome in closely related species of gorillas"

**Supplementary Figures‌**

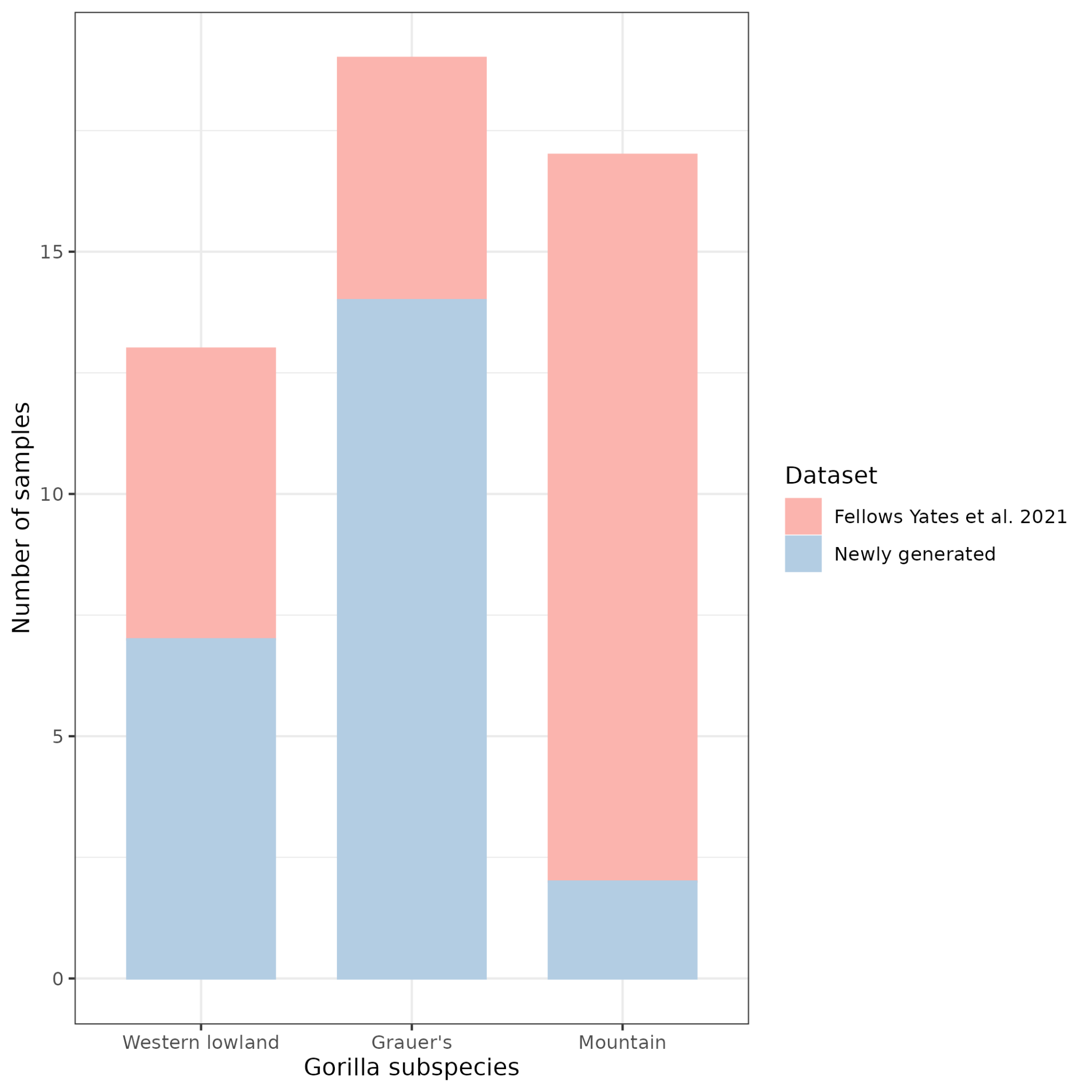

**Figure S1.** Visual summary of the contribution of newly generated and published (Fellows Yates et al. 2021) data to the final dataset.

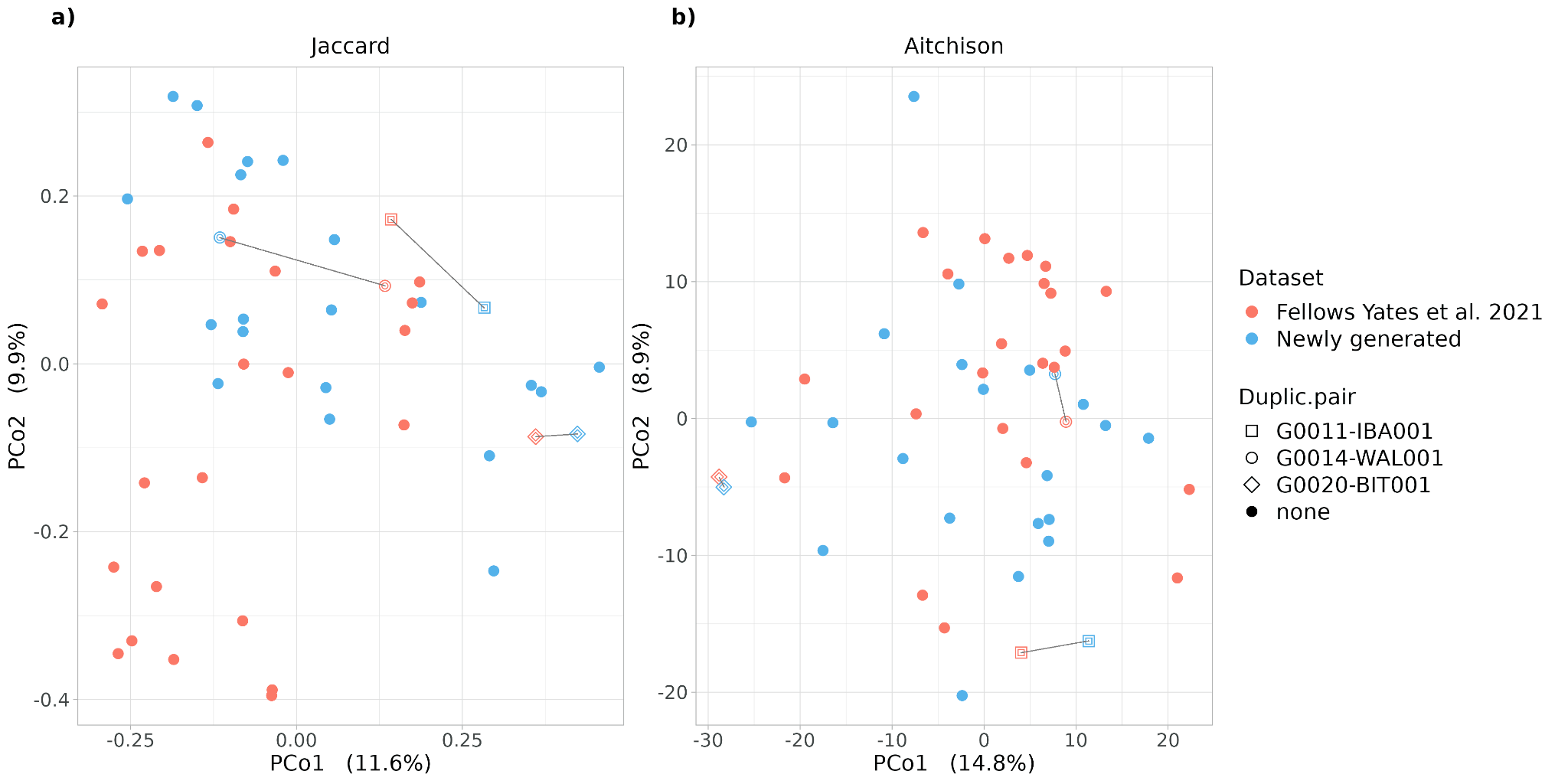
**Figure S2.** Principal coordinate analysis (PCoA) plots based on a) Jaccard distance and b) Aitchison distance highlighting pairs of samples that correspond to the same museum specimen but were processed separately in this study and by Fellows Yates et al. (2021). Duplicate pairs are shown as diamonds and are linked with a grey line. Datasets are displayed in different colours.

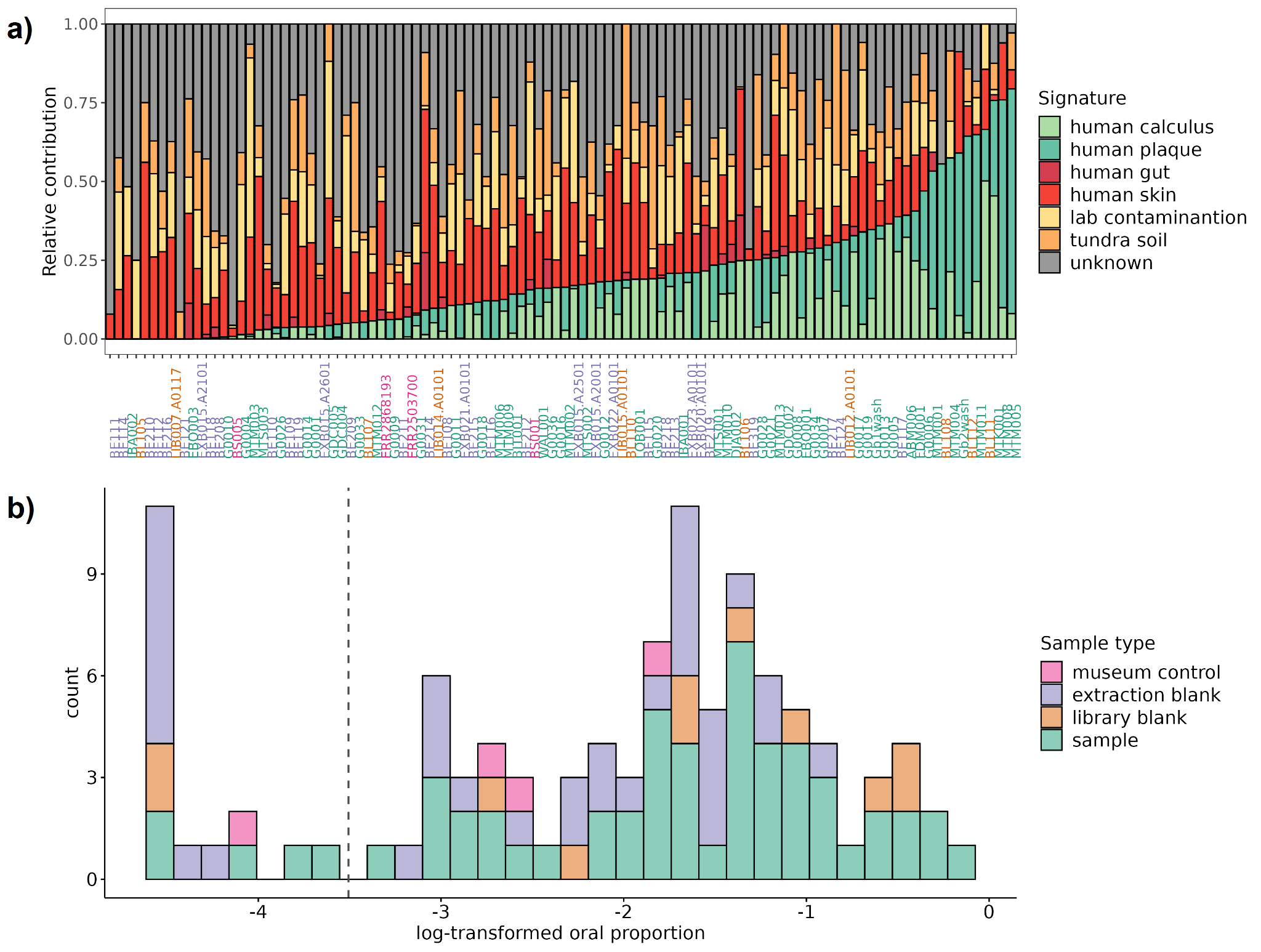

**Figure S3.** (a) Output of FEAST showing the composition of dental calculus-derived microbial communities partitioned into different putatively contributing environments for each sample. Sample ID colours correspond to sample types, as detailed in b). (b) Histogram showing the log-transformed oral proportion (human calculus and human plaque considered jointly) per sample. Only dental calculus samples with more than 300,000 reads are shown here (see Figure S11). Dental calculus samples (coloured in green) with an oral microbiome proportion below 3% (those left of the vertical dashed line) were excluded from analyses. In contrast, all negative controls and museum environment samples were retained. The bars are coloured by sample type: museum controls (pink), extraction blanks (purple), library preparation blanks (orange), specimen samples (green).

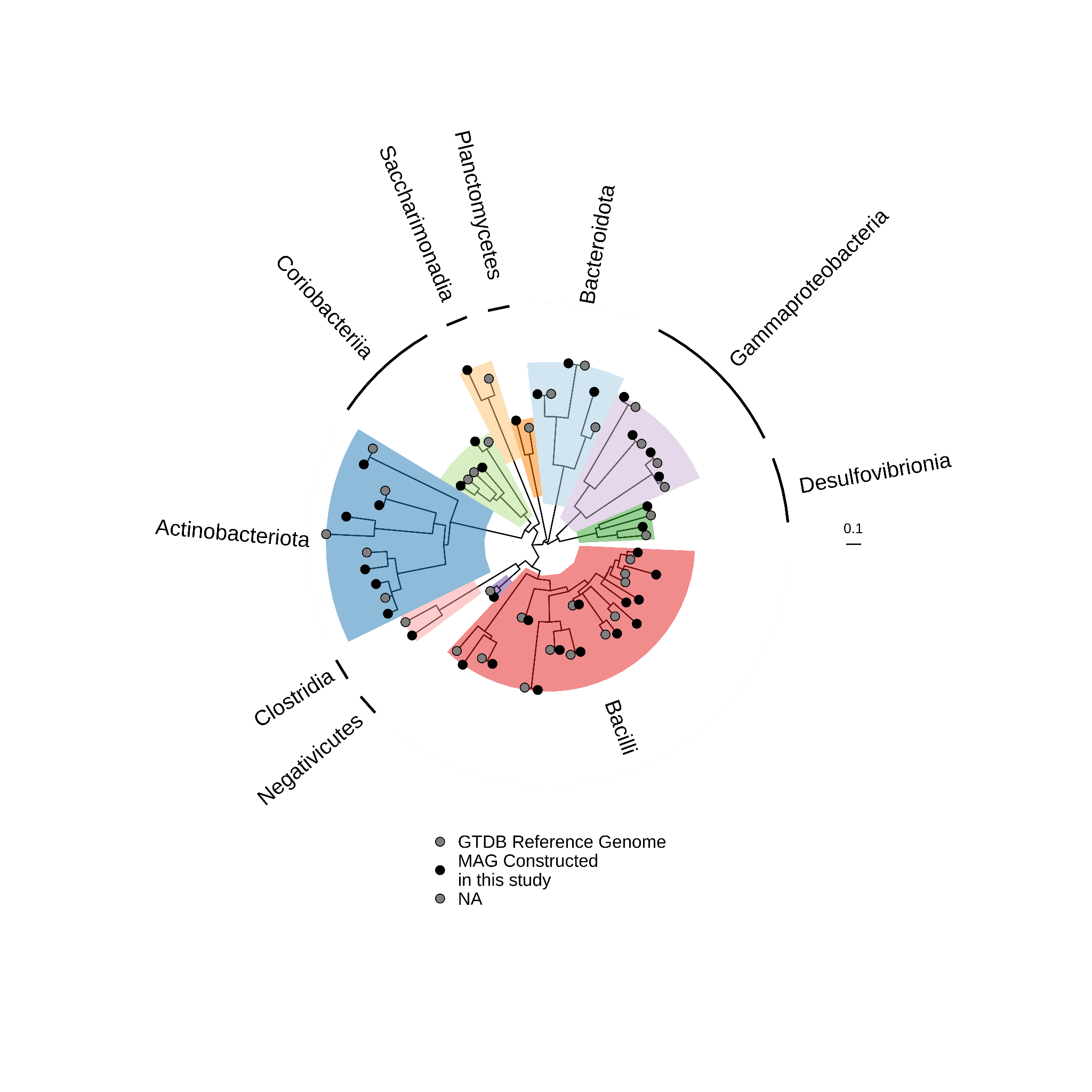

**Figure S4.** MAG phylogeny including the closest related reference genomes available in the taxonomic database, GTDB. *Actinobacteriota*, *Bacilli*, *Bacteriodota* designate distinct bacterial phyla, whereas all others denote bacterial classes. Branch lengths are scaled to the number of substitutions per site.

**Figure S5.** The CLR-normalised abundance of MAGs reconstructed from decontaminated reads. MAG taxonomic identity is provided on the y-axis, and the identity of individual samples is shown on the x-axis. Host subspecies and museum controls are shown on top of the heatmap, highlighted in different colours, including the skull (BS005) and shelf (BS001) swabs, petrous bone (ERR2503700), and skin (ERR286193). MAGs are sorted based on prevalence across all samples, and the absence of a MAG within a sample is shown in grey. Sample G0005 contained many MAGs not found in any other sample, likely due to much deeper sequencing of this sample compared to others (77,307,443 processed reads compared to the mean of 11,365,074 reads per sample). Asterisks denote taxa that are found at higher abundance in at least one museum control compared to any dental calculus samples (red) and taxa likely to be contaminants on the basis of isolation source information (black), as shown in Figure S6.
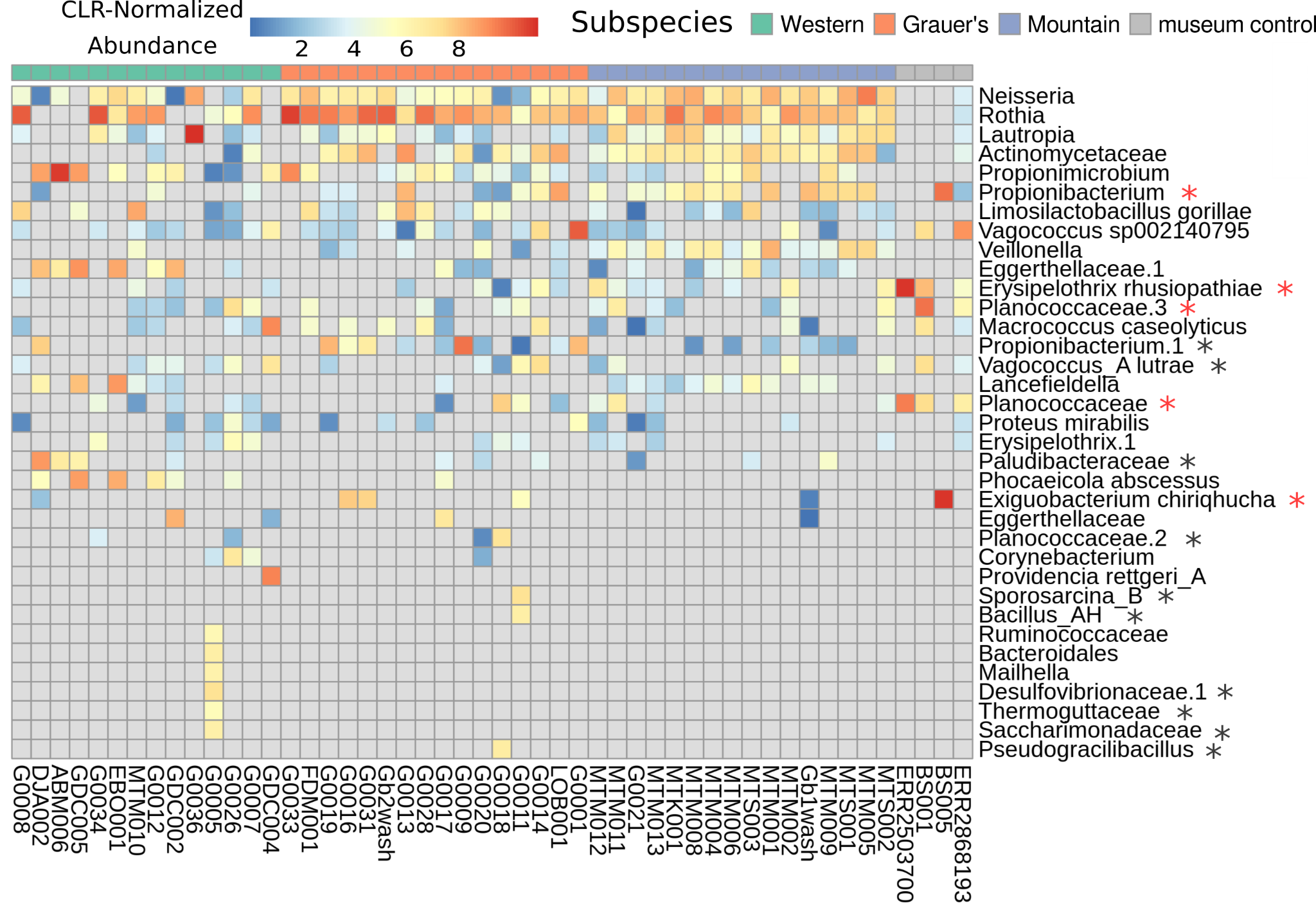

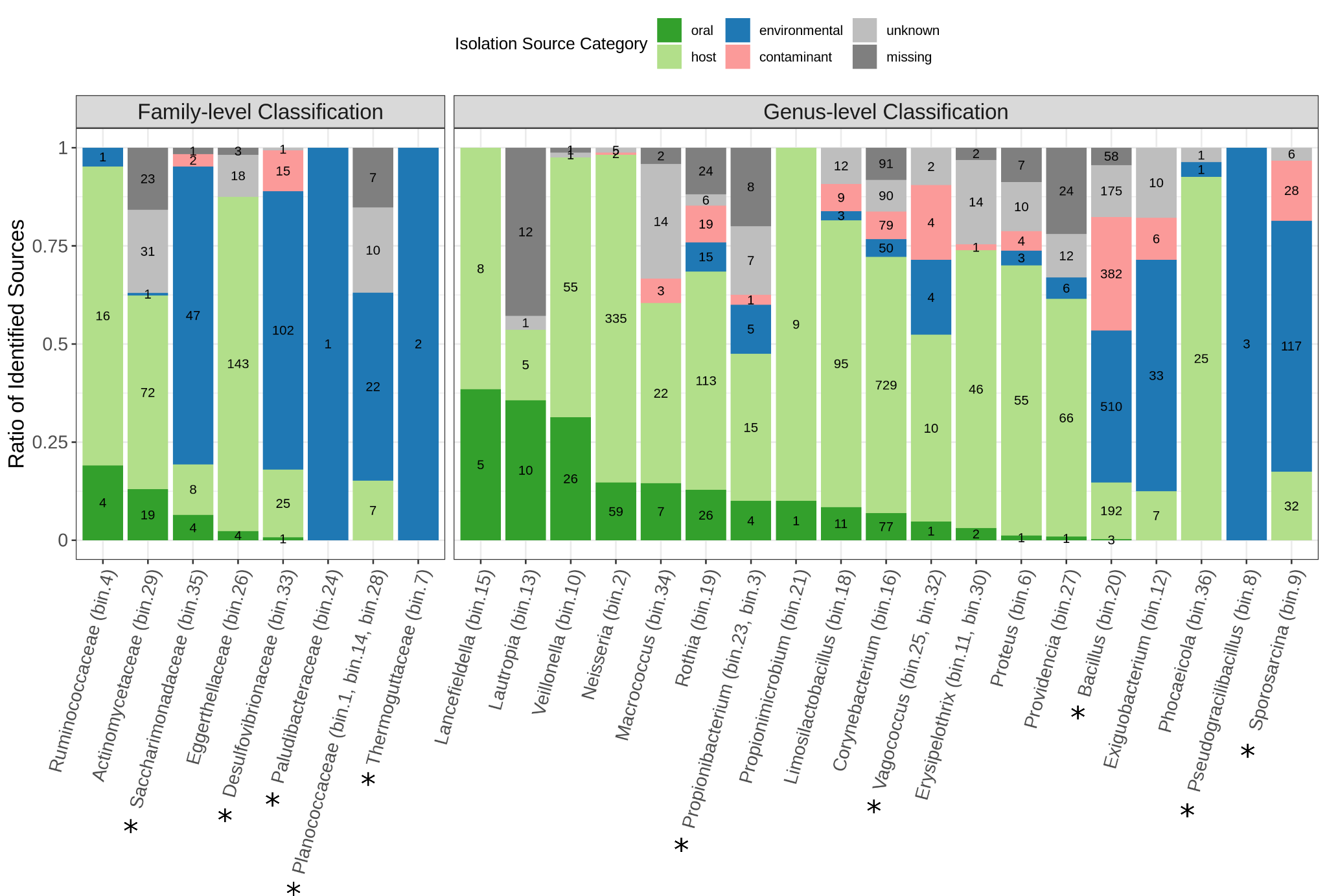

**Figure S6.** Isolation source information for each MAG taxon, as obtained from multiple databases (see Table S8). Broad categories of isolation sources were generated based on the presence of key words. Values within each section of a stacked bar represent the number of unique isolation sources for each category. Taxa with a combined proportion of environmental and contaminate sources greater than 25% were considered as probable contaminants. Asterisks denote MAGs that were excluded on the basis of the overrepresentation in environmental/contaminant isolation sources.

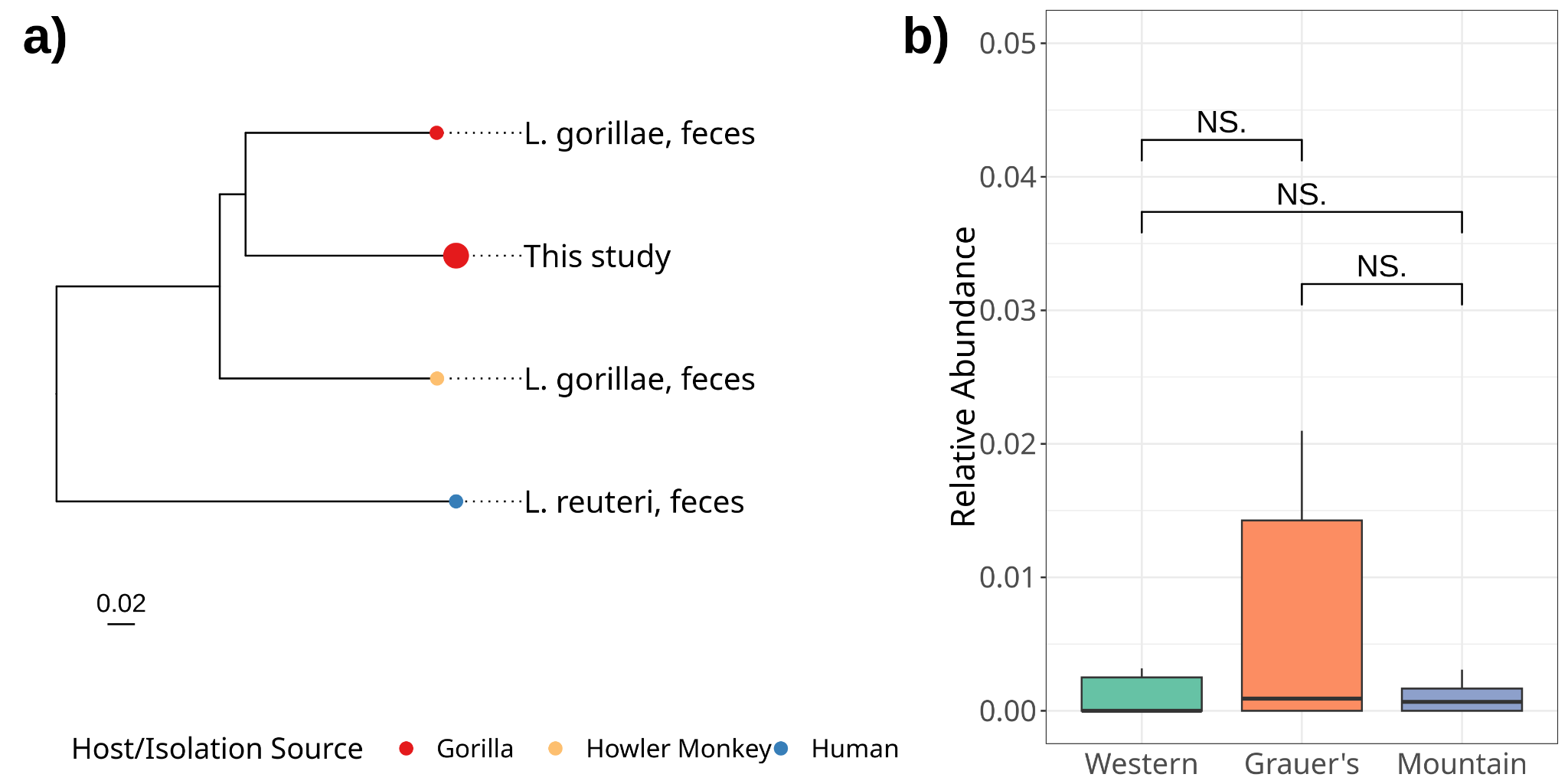

**Figure S7. a)** Maximum-likelihood phylogeny based on alignment of core gene sequences from *Limosilactobacillus gorilla*. Tips include host species identity and the isolation source for each genome. All nodes have complete support (1.0) after 100 bootstrap replicates, and scale bar units are the number of substitutions per site. **b)** The relative abundances (CLR transformed) between different gorilla subspecies in *Limosilactobacillus gorilla*, with the results of a Wilcoxon test are denoted by brackets above boxplots (NS = not significant).

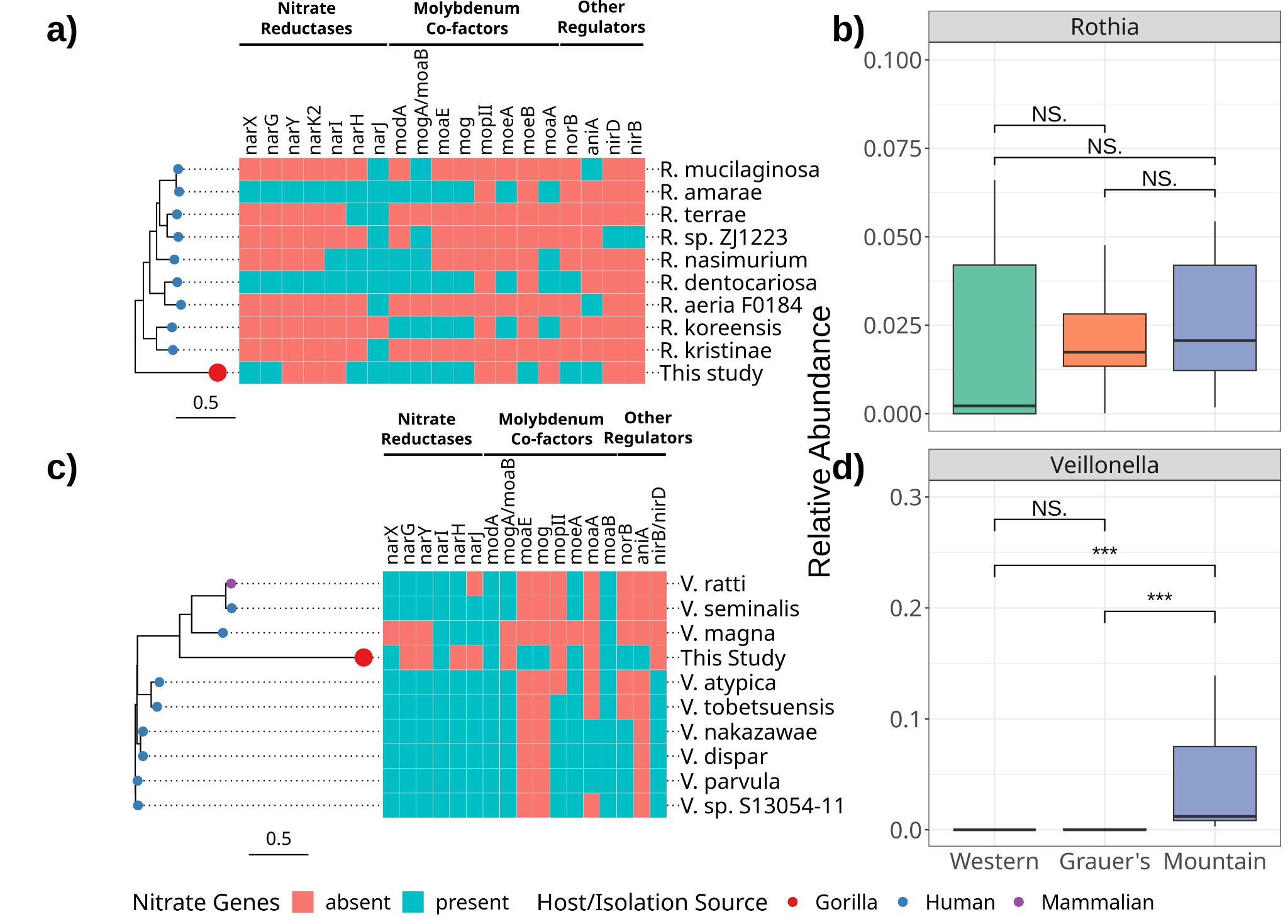

**Figure S8.** Maximum-likelihood phylogenies based on alignment of core gene sequences from MAGs of **a)** *Rothia* and **c)** *Veillonella*, as well as additional published genomes for each genus. Tip points are coloured by host species identity. Scale bar units are the number of substitutions per site. All nodes have support values of 1 after 100 bootstrap replicates. To the right, the presence of key genes involved in the reduction of nitrate in the oral cavity is shown for each genome. Panels **b)** and **d)** show the relative abundances (CLR transformed) of *Rothia* and *Veillonella* MAGs, respectively, in different gorilla subspecies. The results of a Wilcoxon test are denoted by brackets above boxplots (NS = not significant; *** = p > 0.001).

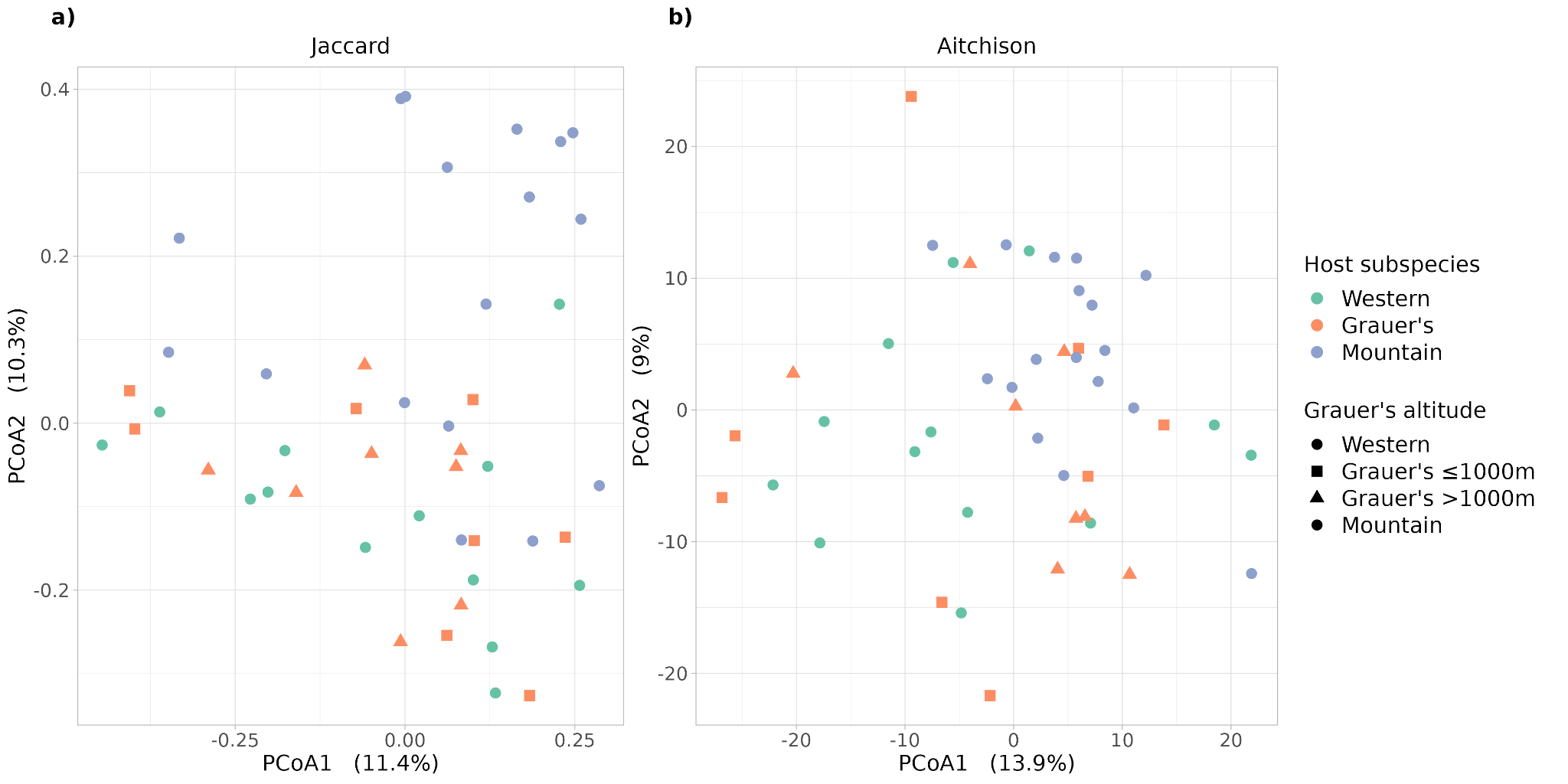

**Figure S9.** Principal coordinate analysis (PCoA) plots highlighting the effect of altitude on the oral microbiomes of Grauer’s gorillas based on a) Jaccard distance and b) Aitchison distance. Host subspecies is displayed in different colours. Grauer’s gorillas from low altitudes are shown as squares and from high altitudes as triangles.

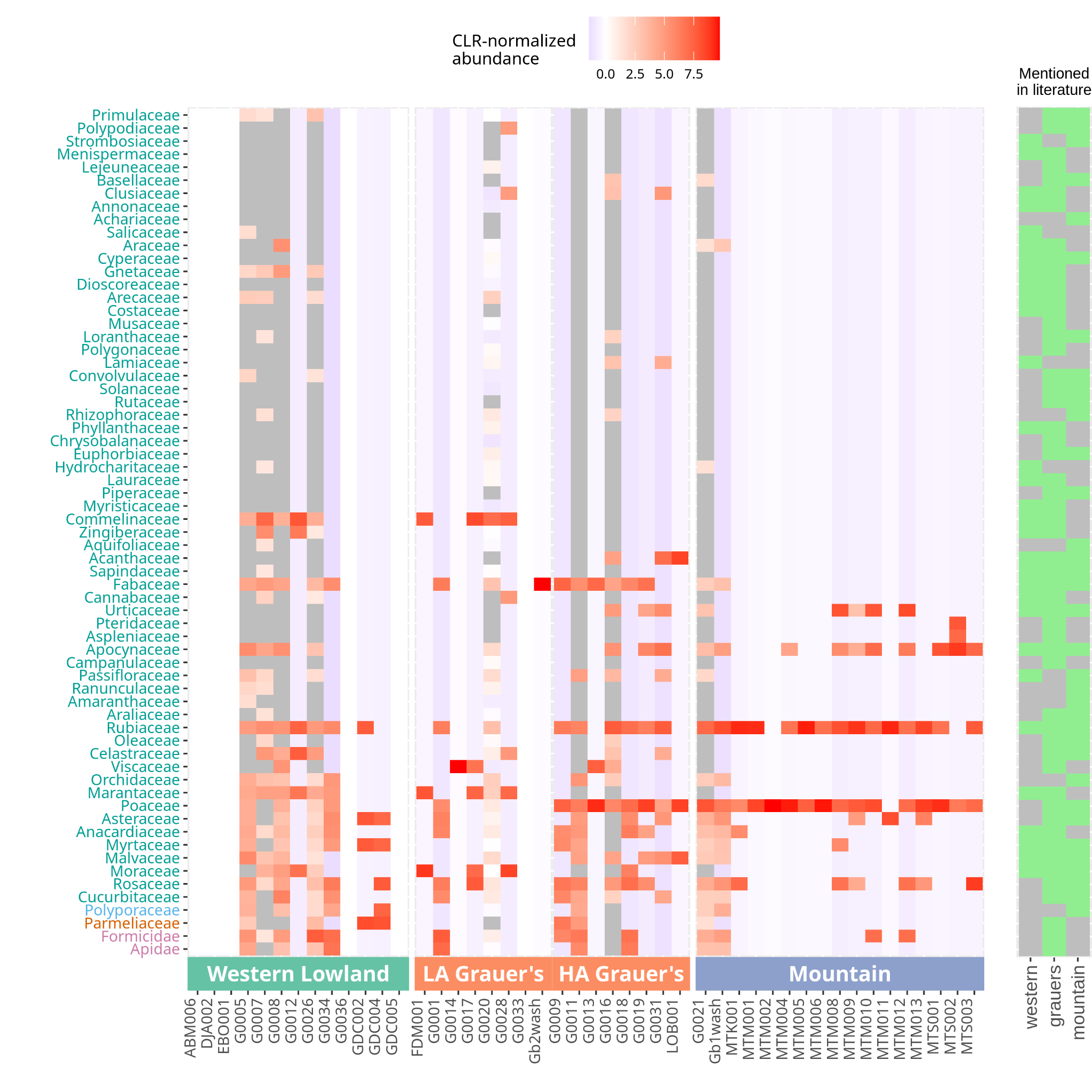

**Figure S10.** Heatmap based on CLR-normalised abundance of eukaryotic families that are known to be consumed by gorillas. Labels of the y axis are coloured based on the phylum (cyan: higher plants; blue: Basidiomycota; orange: Ascomycota; pink: arthropods). The bar on the right indicates if the family has been listed as part of the diet of each gorilla subspecies (grey: absent; green: present).

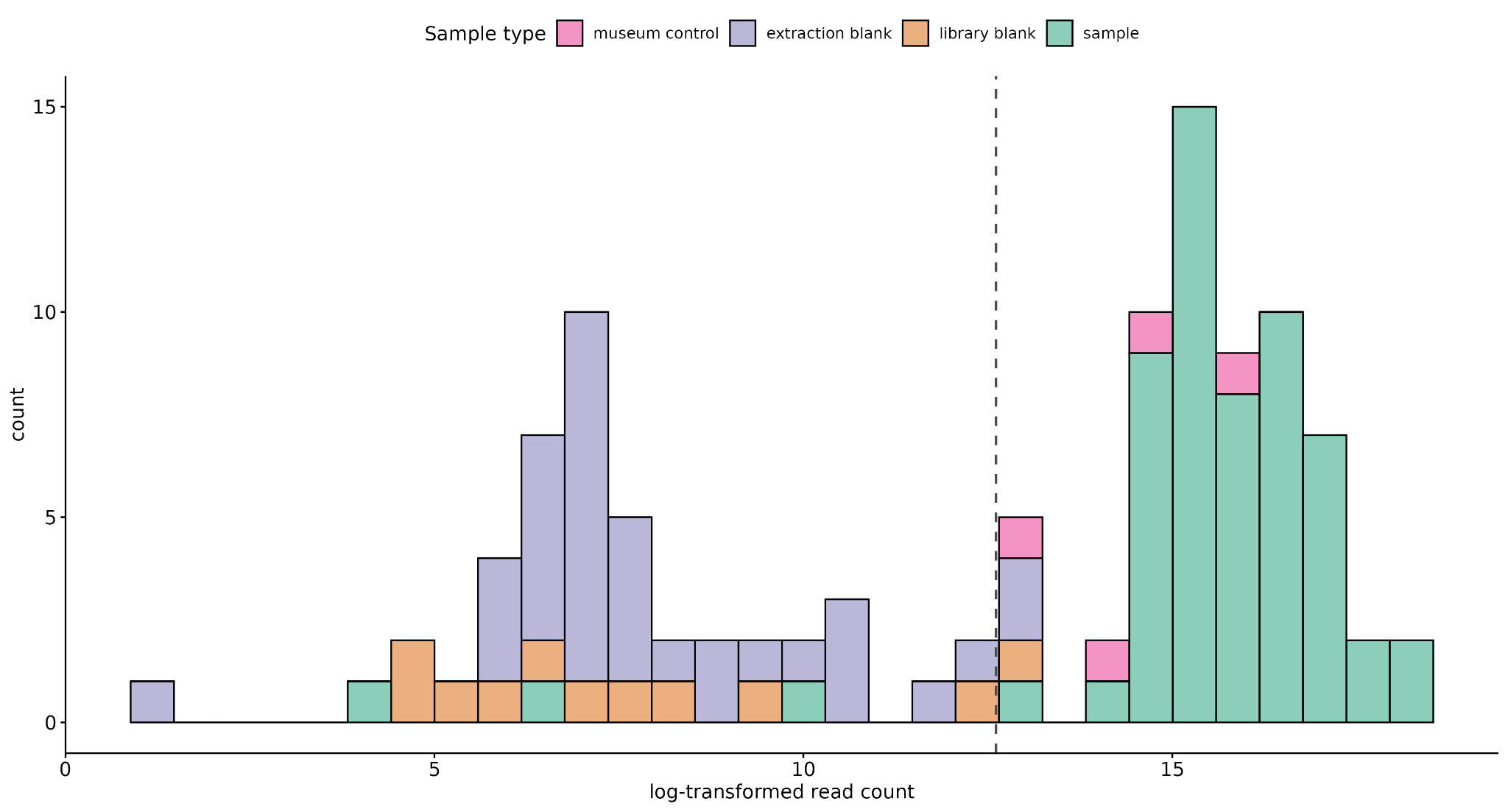

**Figure S11.** Histogram showing the read count (log-transformed) after pre-processing for all samples that were subjected to taxonomic classification. The bars are coloured by sample type: museum controls (pink), extraction blanks (purple), library preparation blanks (orange), specimen samples (green). Dental calculus samples (green) containing less than 300,000 reads (those left of the vertical dashed line) were excluded.

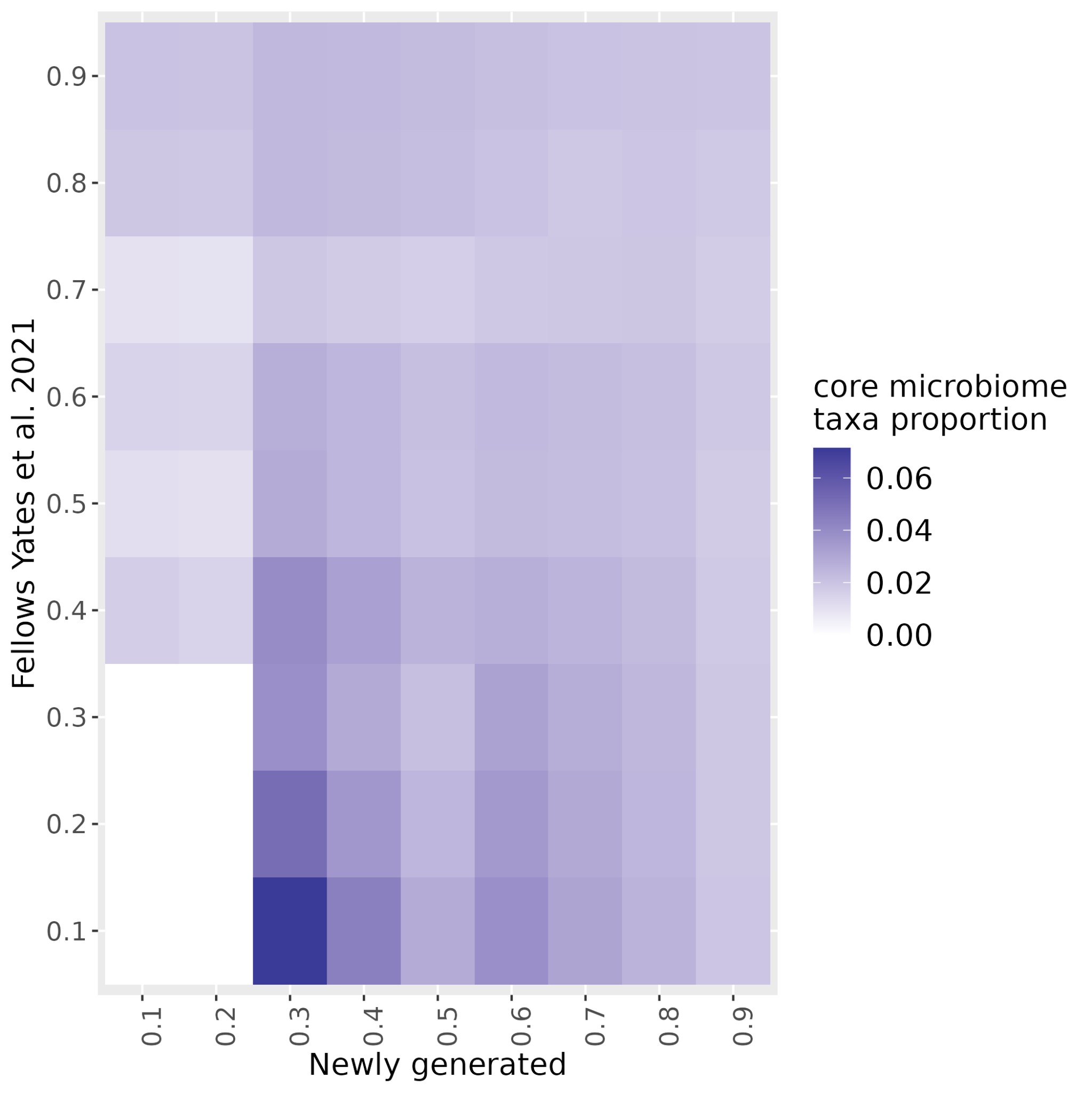

**Figure S12.** Heatmap showing the proportion of oral taxa among the total taxa identified as contaminants with the R packaged decontam using different thresholds for the newly generated (x-axis) and Fellows Yates et al 2021 (y-axis) datasets.
